# The effect of a reduction in population size on mean fitness and inbreeding depression

**DOI:** 10.64898/2026.05.15.725556

**Authors:** Eugenio López-Cortegano, Brian Charlesworth

**Affiliations:** Institute of Ecology and Evolution, School of Biological Sciences, University of Edinburgh, Edinburgh EH9 3FL, United Kingdom; Aviagen Ltd., Newbridge, Edinburgh EH28 8SZ, United Kingdom

**Keywords:** genetic load, inbreeding load, inbreeding, inbreeding depression, genetic purging, mean fitness, conservation genetics

## Abstract

A sudden reduction in population size increases the rate of genetic drift, reducing variability and increasing the mean level of homozygosity. The resulting increase in exposure of strongly deleterious, recessive or partially recessive alleles to selection against homozygotes may lead to their being purged from the population, potentially allowing mean fitness to increase after an initial decline, and accelerating the decline in inbreeding depression associated with reduced variability. However, detailed population genetic theory on the effects of population bottlenecks on mean fitness and inbreeding depression remains limited. We develop a theoretical framework for small, randomly mating populations founded from a large population at mutation–selection–drift equilibrium, using both simulations and approximate analytical results. These provide quantitative predictions for the dynamics of the population’s mean fitness and level of inbreeding depression following a bottleneck. In particular, we derive an approximate expression for the time needed for mean fitness to recover after an initial decline; such a recovery requires selection to be sufficiently strong relative to drift, and mutations to be sufficiently recessive. In contrast, weakly deleterious mutations cause reductions in mean fitness and inbreeding depression that are similar in size to those predicted from increases in homozygosity for neutral alleles, over the relatively short timescales considered here.

## Introduction

Fitness decline due to genetic drift affecting deleterious mutations is likely to be a major threat to the survival of small, endangered populations (Lande 1994; Latter 1998; Hedrick and Kalinowski 2000; Crow 2008; Frankham 2010). Increases in both the mean frequencies and levels of homozygosity of deleterious mutations that typically segregate at low frequencies in natural populations contribute to this phenomenon (Wright 1931; Charlesworth 2018; Kardos et al. 2025). The effect of mean homozygosity on fitness in small populations depends in part on the inbreeding load (*B*), the measure of the rate at which the natural logarithm of fitness declines with an increase in the inbreeding coefficient *F* (Morton et al. 1956), which itself depends on the frequencies and fitness effects of deleterious alleles. There has consequently been much interest in estimating *B* for the purpose of assessing population health and guiding conservation strategies. Empirical evidence suggests that *B* is often substantial in wild populations (O’Grady et al. 2006, Nietlisbach et al. 2019, Stoffel et al. 2021), leading to the notion of a minimum viable population size below which there is a high risk of extinction (Jamieson and Allendorf 2012; Frankham et al. 2014a, b; Franklin et al. 2014; Caballero et al. 2017; Pérez-Pereira et al. 2022).

However, the increased frequency of homozygotes for recessive or partially recessive deleterious mutations in small, randomly mating populations enhances the ability of natural selection to reduce their frequencies (Crow 1970; Hedrick 1994), potentially mitigating the loss of mean fitness caused by their increased level of homozygosity. As a consequence of this increased rate of elimination of rare deleterious alleles, small populations may also experience a reduction in *B* in addition to the effect on *B* of reduced variability due to drift. This enhanced effectiveness of selection against deleterious mutations is known as “genetic purging”, and may lead to a recovery of mean fitness after an initial period of inbreeding depression (Wang et al. 1999; García-Dorado 2012; Charlesworth 2018). Understanding the consequences of purging for mitigating the effects of genetic drift in small populations is fundamental to an understanding of their evolution and the survivability of endangered species.

Genetic purging remains less understood than inbreeding depression itself, which is usually studied under conditions that avoid changes in allele frequencies (Glémin 2003, Hedrick and García-Dorado 2016). Theoretical models suggest that purging primarily removes strongly deleterious and highly recessive alleles from small populations, whereas weakly selected mutations eventually drift to fixation (Kimura et al. 1963, Wang et al. 1999, Glémin 2003). Recent genomic studies support this distinction, showing that small populations tend to accumulate mutations with small fitness effects, whereas strongly deleterious variants tend to decline in frequency due to purifying selection (e.g., Xue et al. 2015, Kardos et al. 2016, Robinson et al. 2016, Grossen et al. 2020, Stoffel et al. 2021, Kleinman-Ruiz et al. 2022, Kyriazis et al. 2023a, Lavanchy et al. 2024, Stuart et al. 2025). Some genomic studies have also suggested that purging may cause a reduction in inbreeding depression in certain populations (Campos et al. 2010, Robinson et al. 2022, Kyriazis et al. 2023a). However, there is limited knowledge regarding the conditions and timescale required for the purging of deleterious mutations, and the dynamics of any associated recovery of mean fitness. Without a rigorous theoretical framework, it is hard to determine whether failure to detect purging in field and experimental studies comes from insufficient data or is a real phenomenon.

This study uses computer simulations and analytical theory to quantify the dynamics of the effects of a population bottleneck on the frequencies of deleterious alleles, population mean fitness and level of inbreeding depression. In particular, we derive the conditions under which purging is expected to occur in a randomly mating population. We show that it requires sufficiently strong selection relative to genetic drift (*N*_*e*_*s* ≫ 1) and a sufficiently low dominance coefficient (*h* < ⅓), where *N*_*e*_ is the post-bottleneck effective population size, *s* is the selection coefficient against mutant homozygotes and *hs* is the reduction in fitness for mutant heterozygotes. Based on these conditions, we develop analytical predictions under two alternative selection regimes, corresponding to strongly (*N*_*e*_*s* ≫ 1) and weakly (*N*_*e*_*s* ≲ 1) selected deleterious mutations. For each regime, we use classical approximations to the selection equations and diffusion equation theory to predict changes in mean allele frequencies and fixation indices (inbreeding coefficients). These are then used to derive the expected changes in genetic (*L*) and inbreeding (*B*) loads over time. Under strong selection, we show that the timescale over which purging becomes manifest as a recovery of mean fitness spans a few tens of generations for a wide range of small population sizes and mutational effects. The extent of fitness recovery, however, depends strongly on the genetic architecture and the severity of the bottleneck, with smaller populations experiencing more rapid purging but at a higher cost of fixation of deleterious alleles. Under some parameter combinations, the mean fitness of a bottlenecked population can eventually exceed that of its large ancestral population. In contrast, more weakly selected and less recessive mutations produce changes in mean fitness and inbreeding depression that are close to those expected from neutral increases in homozygosity.

### A model of the effects of selection and drift in a bottlenecked population

#### The model of selection against deleterious mutations

We use the standard model of a diallelic autosomal locus in a randomly mating population, with alleles A_1_ (wild-type) and A_2_ (mutant) that are subject to mutation at rate *u* from A_1_ to A_2_. Reverse mutations from A_2_ to A_1_ are neglected, because of the short timescale under consideration, despite their being necessary for the maintenance of a stationary distribution of allele frequencies in the ancestral population (Wright 1931). At a given locus, the frequencies of A_1_ and A_2_ in an arbitrary generation are *p* and *q* = 1 – *p*, respectively. Following standard notation, the fitnesses of A_1_A_1_, A_1_A_2_ and A_2_A_2_ are 1, 1 – *hs* and 1 – *s*, respectively, where *s* is the selection coefficient and *h* is the dominance coefficient.

We consider a large set of independent autosomal loci, each subject to identical selection and mutation parameters. The probability distribution of the deleterious allele frequencies generated by mutation, selection and drift represents the states of these loci, and determines population-level statistics such as the mean allele frequency, the genetic load and the inbreeding load. This distribution can be regarded as representing the state of a large number of independent loci within the same population. We first describe these statistics for the ancestral population and then analyse their behavior following a population bottleneck.

#### Population genetic statistics for the ancestral population

We first consider a large ancestral population with effective size *N*_0_, assumed to have remained sufficiently large (*N*_0_*s* ≫ 1) for deleterious mutations to reach mutation-selection-drift equilibrium. In this population, mutation continuously introduces deleterious alleles, selection opposes their increase in frequency, and genetic drift generates stochastic variation in allele frequencies among loci. For mutations that are not completely recessive and that follow the condition *N*_0_*hs* ≫ 1 so that selection dominates over drift, the stationary probability distribution is a gamma distribution with mean *q*^∗^ ≈ *u /* (*hs*) ≪ 1, which is the same as the deterministic value (Haldane 1937), and has an approximate variance *q*^∗^/ (4*N*_*o*_*hs*) (Nei 1968). For completely recessive mutations (*h* = 0), with 4*N*_*o*_ ≪ 1 (as is normally the case) and 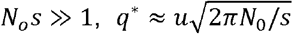, with a variance of approximately *u*/*s* (Nei 1968).

The equilibrium distribution of allele frequencies determines the expectations of the genetic and inbreeding loads, whose values for non-recessive mutations depend primarily on the mutation rate when the above conditions are met, and are approximately the same as the infinite population expressions of Haldane (1937), Morton *et al*. (1956) and Charlesworth and Charlesworth (1987). The expected value of the genetic load for a single locus in the ancestral population, defined as the reduction in the population mean fitness relative to a value of 1 for A_1_A_1_ homozygotes (Crow 1970), is 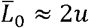 for mutations with *h* bounded away from zero; for completely recessive mutations, 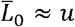 (Haldane 1937; Nei 1968).

The inbreeding load with respect to a single locus, *B*, is defined as the difference between the mean fitness of a randomly mating population and the mean fitness of a fully inbred population with the same allele frequency (Morton et al. 1956). For a locus with *q*\* ≪ 1 and *h* ≫ *q*\*, the expected value of *B* for the ancestral population is 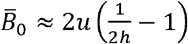 (Charlesworth and Charlesworth 1987, Equation 3b). For completely recessive alleles with 4*N*_*o*_*u* ≪1, Equations (15) of Nei (1968) imply that 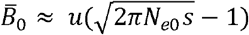.

#### Population genetic statistics for a bottlenecked population

We next assume that the population has undergone a bottleneck, reducing its size to *N* breeding individuals, where *N* ≪ *N*_0_. We assume a Wright-Fisher population model, so that the effective size (*N*_*e*_) of the bottlenecked population equals *N*. Relative to the ancestral population, the smaller effective population size increases the influence of genetic drift, producing greater stochastic fluctuations in allele frequencies. This results in an increase over time in the variance in allele frequencies and in the mean frequency of homozygotes, although selection continues to remove deleterious alleles whenever it is sufficiently strong relative to drift (*Ns* ≫ 1).

When a bottlenecked population is founded by sampling *N* breeding individuals from a large source population with an expected frequency of the deleterious allele of *q*\*, the initial population has a net probability of approximately *P* = 1– 2*Nq*\* of containing no copies of A_2_ at a given locus and 1 – *P* = 2*Nq*\* that it contains at least one copy, provided that 2*Nq*\* ≪ 1, so that second- and higher-order terms in 2*Nq*\* can be ignored (Figure 1). This approximation is justified by the fact that the parameter sets used here are such that *q*\* < 5 × 10^−6^ and *N* ≤ 500, so that 2*Nq*\* < 5 × 10^−3^. The first case contributes nothing to population genetic statistics dependent on non-zero *q*, except for contributions from mutations that occur after the bottleneck. In the second case, the expected initial frequency of A_2_ is *q*_0_ = 1/(2*N*). This is, of course, far higher than *q*\*.

**Figure 1.**
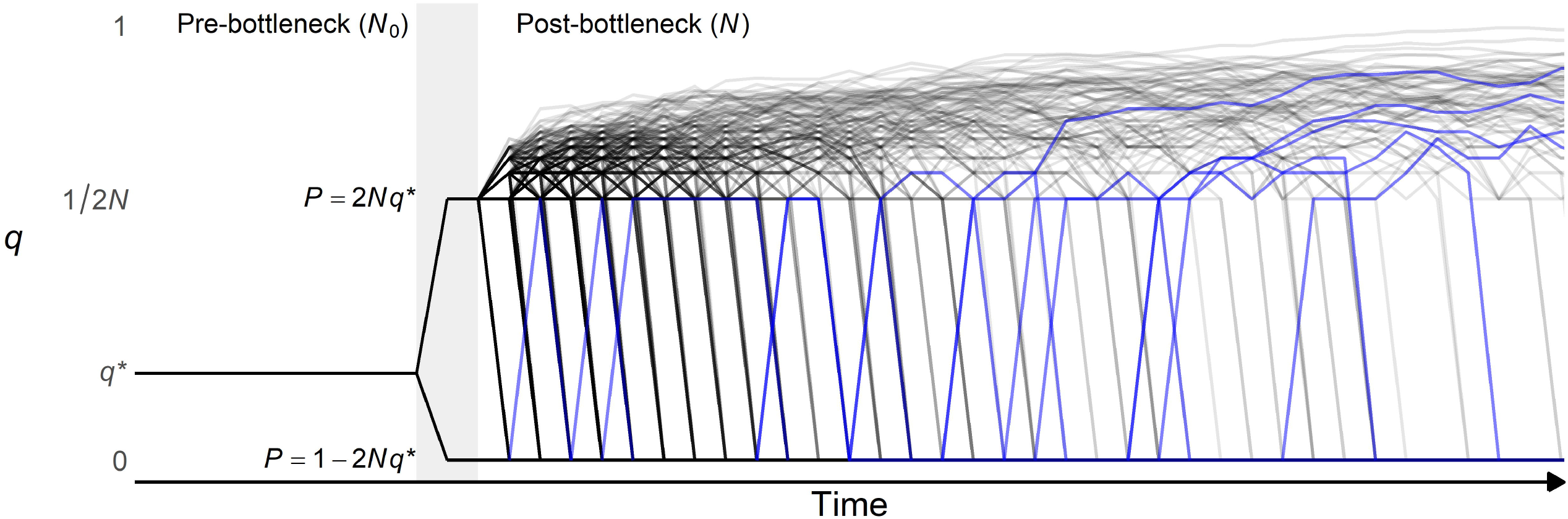
Allele frequency dynamics following a bottleneck. After a bottleneck (represented by the background grey bar), allele frequencies in a sample of *N* individuals derived from a large population of size *N*_0_ change from their values at mutation-selection-drift equilibrium (mean frequency = *q*\* ≪ 1) to either zero (with approximate probability 1-2*Nq*\*) or 1/2*N* (with approximate probability 2*Nq*\*) in the new population, assuming that *N* ≪ *N*_0_. Following the bottleneck, drift and selection are the dominant forces driving the frequencies of both initially segregating alleles (in grey) and new mutations arising post-bottleneck (in blue), which become either fixed or lost over time. For the purposes of illustration, the Y axis is plotted on a log_10_ scale.

In general, there will be variation among independent evolutionary replicates in the probability that the founder population has a copy of A_2_, because allele frequencies in the ancestral population follow a probability distribution around *q\** (see above). Use of a fixed probability of 2*Nq\** thus underestimates the variability among replicates in whether A_2_ survives the bottleneck. However, the overall probability that A_2_ is present is still 2*Nq\**. It follows that the use of a fixed value of 2*Nq\** has no effect on the means of the population statistics, since the initial frequency of A_2_ is fixed at 1/(2*N*).

In the absence of any mutational contributions to allele frequency change, the expected frequency of A_2_ at segregating loci must therefore start to decline over the generations, provided that selection is sufficiently strong in relation to drift, regardless of the level of dominance. In this sense, purging of deleterious mutations at segregating loci inevitably occurs over the short term, although the overall expected value of *q*,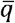 (which includes both segregating and fixed loci) is unchanged at *q*\* in the first generation of the bottleneck. The net change in 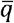 over time depends, however, on events at both initially segregating and initially non-segregating loci, including mutational contributions, and cannot be predicted intuitively. Eventually, the accumulation of new mutations at both types of loci (including reverse mutations from A_2_ to A_1_) will cause the system to reach a new equilibrium under mutation, selection and drift (e.g., Lande 1998).

The realised fixation index for loci with selection coefficient *s, F*_*s*_, is defined as the ratio of the variance in allele frequency at these loci to the product 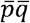, where overbars indicate expected values taken over the entire probability distribution of *q*, including loci at which A_2_ alleles were initially lost; we also have 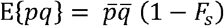 (e.g., Wright 1951). We use the term “fixation index” rather than “inbreeding coefficient”, as we are concerned with deviations from Hardy-Weinberg frequencies rather than identity by descent.

The expected allele frequency and fixation index together determine the expected genotype frequencies, and hence the expected mean fitness, which for a single locus is given by 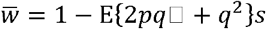. If selection is sufficiently strong in relation to mutation and drift in the ancestral population, it is reasonable to assume that 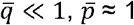. This implies that 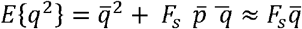 and 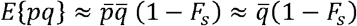, if terms of order 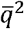 are neglected. Under this rare-allele assumption, the expected genetic load, 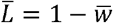, is given approximately by:

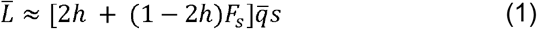

Similarly, the inbreeding load *B* for a single locus is defined as the difference between the mean fitness of a randomly mating population, 1 − [2*h* + (1 − 2*h*)*q*]*qs*, and the smean fitness of a fully inbred population with the same allele frequency, 1 − *qs*, so that *B* = (1 − 2*h*)*pqs* (Morton et al. 1956). The expected value of *B* over the distribution of allele frequencies is thus:

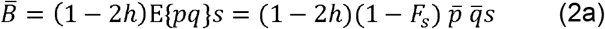

Using the rare-allele assumption described above, this expression can be approximated by:

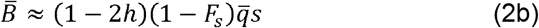

Inspection of Equations (1) and (2) brings out several important points about the probable effect of a bottleneck on the load statistics. First, a decrease in 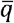 due to purging of deleterious alleles tends to reduce 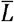, whereas an increase in *F*_*s*_ due to drift has the oppositeeffect. It is therefore unclear *a priori* how the bottleneck will affect the mean fitness of the population. Second, an increase in and a reduction *F*_*s*_ in 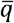 over time will both act to reduce 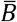, so that we can always expect the inbreeding load to decrease. Third, it is likely that *F*_*s*_ will increase more slowly over time than the fixation index *F* for neutral loci, due to selection opposing drift by pushing allele frequencies towards their equilibrium values. Because of the *F*_*s*_ dual effects of and 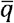 on 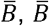 could either decrease faster or more slowly than what is expected from the trajectory of the neutral *F* over time. A faster observed decrease in 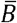 than predicted from neutral *F* would suggest purging of deleterious alleles.

## Methods of analysis

### Changes in allele frequencies following the bottleneck

The general expression for the change in the frequency of A_2_ due to selection and mutation at a locus with allele frequency *q* in a given generation is:

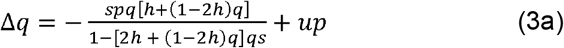

(see Crow and Kimura, 1970, p.183). Note that the allele frequencies and other population genetic statistics are functions of the time *t* since the bottleneck; to avoid unnecessary notation, we represent them as explicit functions of *t* only when it is essential to display their time dependence.

If terms in (*qs*)^2^ can be neglected relative to *qs* when *h* > 0, or terms in *q*^3^*s*^2^ relative to *q*^2^*s* when *h* = 0, this expression reduces to:

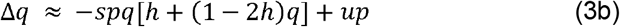

This expression is likely to be accurate for allele frequencies close to zero, even with large *s* values, and so should be valid with strong selection over the short timescales considered here. The net approximate change in the expectation of *q*, after approximating 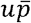 by u, is given by:

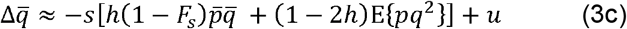

Equations (3) provide the basis on which results for the time course of changes in the expected genetic and inbreeding loads can be studied, either by various types of approximation or by computer simulations of single loci. Table 1 below provides an overview of the analytical methods and corresponding equations used throughout the paper.

**Table 1.**
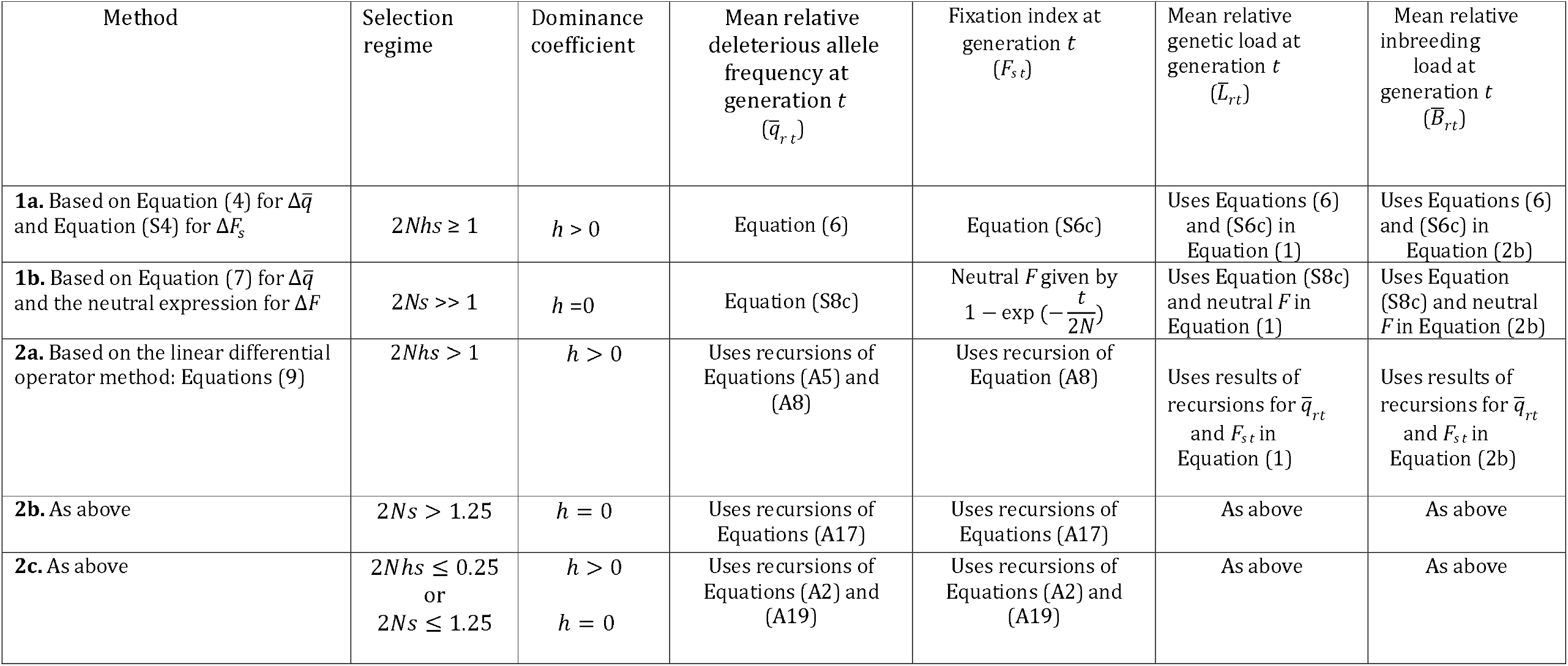
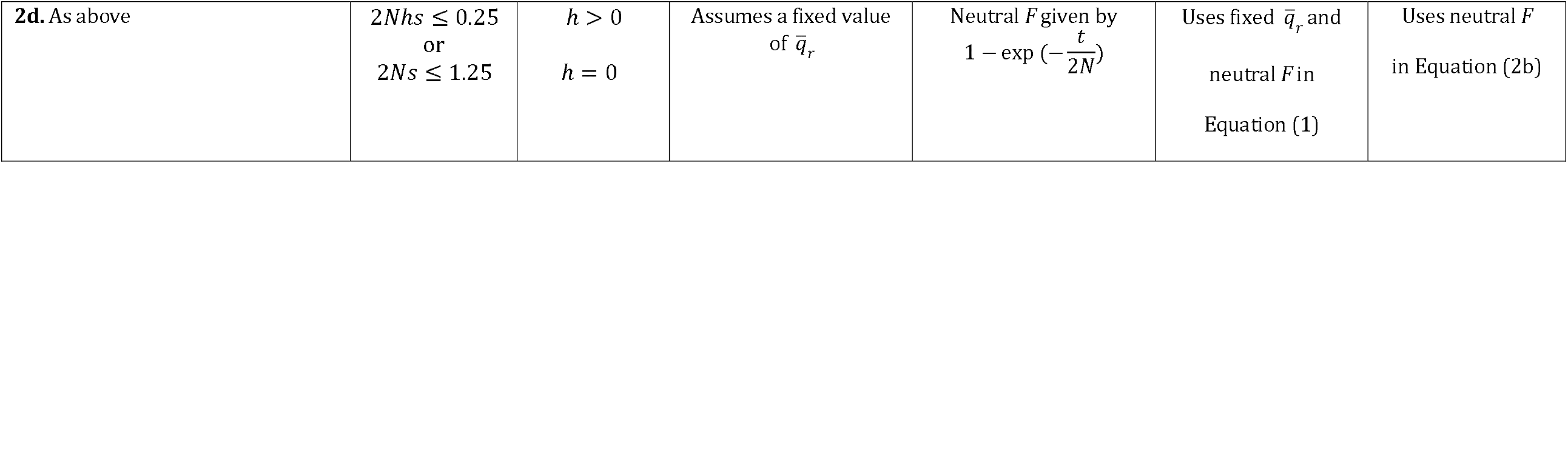
Summary of the main analytical methods and the corresponding equations used to predict the population genetic statistics. For each method, the applicable equations are listed according to the corresponding selection regime (in terms of 2*Nhs* or 2*Ns*) and dominance coefficient (*h*). Note that the mean deleterious allele frequency and the mean genetic and inbreeding loads referred to here, and displayed in the figures, are all expressed relative to their values in the ancestral population, as denoted by the subscript *r*.

| Method | Selection regime | Dominance coefficient | Mean relative deleterious allele frequency at generation *t*<br>($\bar{q}_{r t}$) | Fixation index at generation *t*<br>(*F*<sub>*s t*</sub>) | Mean relative genetic load at generation *t*<br>($\bar{L}_{rt}$) | Mean relative inbreeding load at generation *t*<br>($\bar{B}_{rt}$) |
|---|---|---|---|---|---|---|
| **1a.** Based on Equation (4) for Δ$\bar{q}$ and Equation (S4) for Δ*F*<sub>*s*</sub> | 2*Nhs* ≥ 1 | *h* > 0 | Equation (6) | Equation (S6c) | Uses Equations (6) and (S6c) in Equation (1) | Uses Equations (6) and (S6c) in Equation (2b) |
| **1b.** Based on Equation (7) for Δ$\bar{q}$ and the neutral expression for Δ*F* | 2*Ns* >> 1 | *h* =0 | Equation (S8c) | Neutral *F* given by $1-\exp\left(-\frac{t}{2N}\right)$ | Uses Equation (S8c) and neutral *F* in Equation (1) | Uses Equation (S8c) and neutral *F* in Equation (2b) |
| **2a.** Based on the linear differential operator method: Equations (9) | 2*Nhs* > 1 | *h* > 0 | Uses recursions of Equations (A5) and (A8) | Uses recursion of Equation (A8) | Uses results of recursions for $\bar{q}_{rt}$ and *F*<sub>*s t*</sub> in Equation (1) | Uses results of recursions for $\bar{q}_{rt}$ and *F*<sub>*s t*</sub> in Equation (2b) |
| **2b.** As above | 2*Ns* > 1.25 | *h* = 0 | Uses recursions of Equations (A17) | Uses recursions of Equations (A17) | As above | As above |
| **2c.** As above | 2*Nhs* ≤ 0.25<br>or<br>2*Ns* ≤ 1.25 | *h* > 0<br>*h* = 0 | Uses recursions of Equations (A2) and (A19) | Uses recursions of Equations (A2) and (A19) | As above | As above |
| **2d.** As above | 2*Nhs* ≤ 0.25<br>or<br>2*Ns* ≤ 1.25 | *h* > 0<br>*h* = 0 | Assumes a fixed value of $\bar{q}_r$ | Neutral *F* given by $1-\exp\left(-\frac{t}{2N}\right)$ | Uses fixed $\bar{q}_r$ and<br>neutral *F* in<br>Equation (1) | Uses neutral *F*<br>in Equation (2b) |

### A first approximation to the dynamics of population genetic statistics following a bottleneck (Method 1)

Under the assumption of strong selection (*Ns* ≫ 1), we apply the rare-allele approximation and ignore the moments around zero of the distribution of *q* that are higher than the second. This is valid for the first few generations following the bottleneck, since the allele frequencies at the segregating loci are then all close to 1/(2*N*). It should, however, be noted that this procedure is likely to lead to increasing error over time, and we introduce a more exact approach in the next subsection (see Table 1).

This assumption about the higher moments implies that 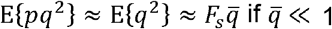; in addition, the rare-allele assumption implies that 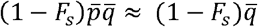. Provided that 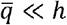, substitution of these expressions into Equation (3c) yields the expression:

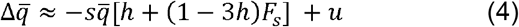

This equation is equivalent to Equation (11) of Whitlock (2002) for 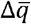 in a structured population, apart from the mutational term, and will be accurate for the period immediately following the bottleneck. Since the mutation and selection terms cancel each other in the ancestral population, so that 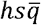 and *u* are very close immediately following the bottleneck (see Equation (5b) below), this equation has the important implication that *h* < ⅓ is required for purging at the level of the overall expected allele frequency.

Method 1a in Table 1 is the approximation for partially recessive deleterious mutations; it is based on Equation (4) by approximating the recursion for 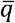 as a differential equation, yielding Equation (6) below. This expression is then combined with the expression for *F*_*s t*,_ the realised fixation index at time *t*, given by Equation (S6c) of Appendix S1. This allows analysis of the dynamics of the expected allele frequency, 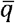, as well as the genetic load, 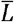, and the inbreeding load, 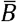(see Equations 1 and 2 for their definitions). We use the convention that *t* = 1 is the generation immediately following the bottleneck, for which *F*_*s*1_ = 1 / (2*N*).*F*_s0_ for the ancestral population can be set to zero as a good approximation, given the assumption that N_0_*hs* ≪ 1

To obtain an expression for 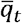, we approximate the discrete recursion equation 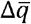 by a differential equation. Equation (4) can be converted to the following differential form for the change in the natural logarithm of 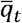:

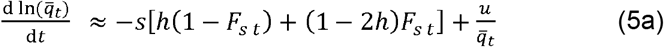

For sufficiently short times after the bottleneck, the mean allele frequency is expected to remain close to its ancestral equilibrium value q^∗^ = *u*/(*hs*) (see the subsection *Population genetic statistics for the ancestral population*), allowing it to be written as 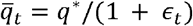, where 0 < *∈*_*t*_ ≪ 1 if purging is occurring. Equation (5a) thus reduces to:

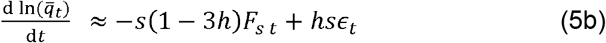

As a first approximation, *∈*_*t*_ is neglected in later calculations (the numerical results displayed in Figure 2 below show that *∈*_*t*_ ≪ 1 over the time periods considered here). Since *∈*_*t*_ >1with purging, neglecting it will overestimate the rate of decline in 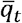, as can be seen in Figure 2.

**Figure 2.**
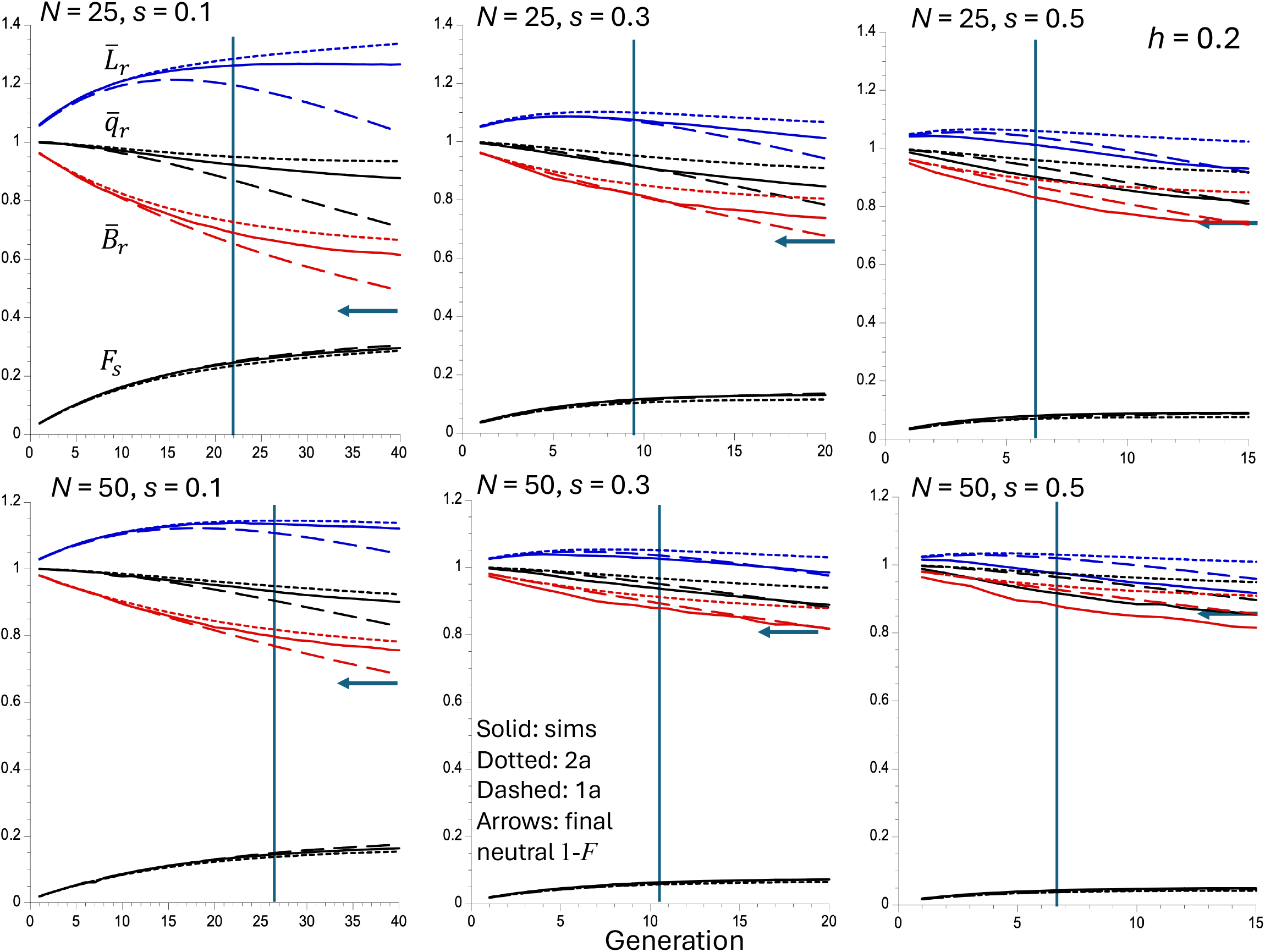
Results of single locus simulations and approximations with *h* = 0.2 and strong selection. The solid curves (in order from top to bottom) in each panel show the means of the following statistics over all replicate simulations: the mean genetic load relative to its equilibrium value 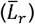, the mean allele frequency relative to its equilibrium value 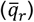, the mean inbreeding depression relative to its equilibrium value 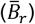 and the mean fixation index at the selected sites (F_s_). The dotted curves are the predictions based on the linear operator Method 2a, and the dashed curves are based on the approximations given by the trajectories for the mean allele frequency using Method 1a and the corresponding expression for the relative values of the genetic and inbreeding loads. The arrows display the final value of one minus the neutral fixation index, 1 – *F*, which corresponds to 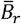 if there is no effect of selection on allele frequencies. The vertical lines indicate the values of the approximate expected time for purging to be manifested (t_c_), obtained from Equation (8). See the *Methods of Analysis* section for details of the simulation method. Note that the mean load eventually declines below its ancestral value 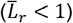 when *s* = 0.5.

It is more convenient for the purpose of displaying results to work with the mean deleterious allele frequency relative to its equilibrium value,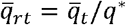. As shown in the Appendix S1 of the Supplementary Material (Equations (S1-S3)), we have:

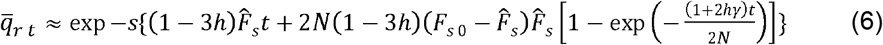

where *γ* = 2*N*_*s*_and *F*_*s t*_ is the fixation index under selection (see Equations S4-S7), with equilibrium value 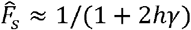 (Equation S5).

The values of the expected genetic load and inbreeding load at time *t*, relative to their values in the initial population,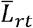 and 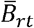, can be obtained by substituting 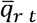 and *F*_*s t*_ into Equations (1) and (2), respectively. The absolute values, 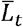 and 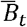, can be found by multiplying the relative values by the ancestral values given by Equations (1) and (2).

For the case of complete recessivity (*h* = 0), Equation (4) is replaced by the following approximation:

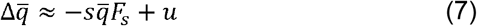

In this case, we have 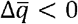 since *F*_*s*_ is larger for the bottlenecked population than its counterpart for the ancestral population.

There is a narrow range of *h* values where *h* is close to 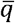 in which neither of these approximations hold, but this can probably be ignored for most purposes given that empirical estimates of *h* are generally well above this range (Di and Lohmueller 2024), even for lethal mutations, as shown by both classical Drosophila studies (Simmons and Crow 1977) and recent molecular evidence from humans (Judd et al. 2026).

Method 1b in Table 1 is the corresponding approximation for the case of completely recessive mutations, based on Equation (7), and is described in the Supplementary Material (Equations (S8-S9)). The mutational contribution is ignored here, as it can be assumed to be negligible when *h* = 0 (Equations (S10)). Additionally, in this case it is necessary to assume that the fixation index *F*_*s*_ is approximated by its value under neutrality (*F*), so that a worse fit to the simulations than with Method 1a is expected, given that *F*_*s*_ < *F*.

### The critical time for purging (t_c_)

A statistic that has recently attracted interest in relation to the effects of population bottlenecks is the minimum expected number of generations required for genetic purging to become manifest at the level of the population mean fitness, *i*.*e*., the “critical time for purging”, *t*_*c*_ (López-Cortegano 2020). We define *t*_*c*_ more precisely as the time after the foundation of the population when the expectation of the mean fitness over the distribution of *q* starts to increase, following an initial decrease due to increased homozygosity for deleterious mutations. Such an increase in 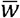 requires *h* < ⅓, the condition for the expected frequency of deleterious mutations to decrease after the bottleneck (see Equation 4).

Under the assumption of strong selection (*Ns* ≫ 1), Appendix S2 of the Supplementary Material provides an approximate expression for the critical time in units of 2*N* generation (Equations (S11-S13)), *T*_*c*_ = *t*_*c*_ /(2*N*). We have:

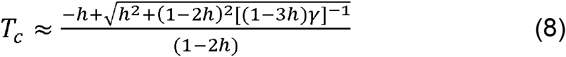

For small *h*, this equation yields 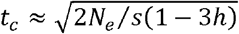. Purging is thus expected to become apparent only after a delay that increases with effective population size and decreases with the strength of selection (see Figure S1A). This implies that smaller populations reach the point of potential fitness recovery faster, but at the cost of higher rates of genetic drift and fixations of deleterious alleles (Figure S1B).

### The linear operator (LO) method for analyzing the population genetics of a bottleneck

A more exact method than those presented so far is provided by the linear differential operator method of Ohta and Kimura (1969), which generates a recursion relation for the expectation of a function of a stochastic variable that obeys the diffusion equation approximation (Method 2 in Table 1). If we use allele frequency *q* as the variable, the general formula for the rate of change per generation in the expectation of *f*(*q*) is:

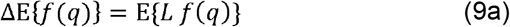

where *L* is a linear differential operator, whose general form is:

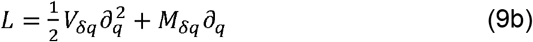

Here, *V*_*δq*_ and *M* _*δq*_ are the variance and the mean of the change in *q* per generation, respectively.

This approach provides a method for obtaining recursion equations for the moments of the probability distribution, which is simpler than an alternative method based on the forward diffusion equation that has recently been used (e.g., Evans et al. 2007; Gravel 2016; Ragsdale 2022). The LO method can be applied to both strong selection, which was modeled above by the more approximate Method 1, and to weak selection. Its implementation in these two situations requires somewhat different methods of approximation, which are described in sections 1 and 2 of the Appendix, respectively (see Table 1). The treatments of strong selection with *h* > 0 (2*Nhs* > 1) and *h* = 0 (2*Ns* > 1) are Methods 2a and 2b, respectively. Method 2c is the more exact LO treatment of weak selection when 2*Nhs* ≤ 1 with *h* > 0 or 2*Ns* ≤ 1 with *h* = 0. Method 2d is a more approximate treatment, which assumes a constant mean allele frequency and a neutral fixation index.

### Computer simulations

To characterize the dynamics of purging on a set of loci with the same selection and dominance coefficients, the accuracy of the approximations used for the single locus dynamics was tested against single-locus simulations, using the Fortran 90 programming language (see Data Availability for the source of the programs and outputs). For each replicate of a simulation with a given value of *N*, a random number *r* was drawn from a uniform distribution on the interval [0, 1) and used to determine whether the deleterious allele A_2_ was initially present in the initial founder population by comparing *r* with 2*Nq\**, where *q\** is the expected frequency of A_2_ for the assigned values of *h, s* and *u*, assuming that 2*N*_0_*S*» 1 for the ancestral population.

As described in the subsection *Population genetic statistics for the ancestral population*, we set *q* = u* / *hs* for *h* > 0 and 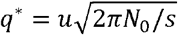 for *h* = 0. If *r* ≤ 2*Nq\**, the initial frequency *q* was set to 1/(2*N*), and the exact Equation (3a) was used to calculate *q* after selection and mutation in a given generation, followed by binomial sampling to generate the new value of *q*. A value of *N*_0_ = 1000 was used for all calculations and simulations with *h* = 0. This process was repeated every generation until A_2_ was fixed or lost, or until the simulation reached a predefined time limit. Otherwise, if A_2_ was initially absent from the population (*r* > 2*Nq\**), new mutations were introduced stochastically each subsequent generation by comparing a new random number with 2*Nu*. If this number was less than 2*Nu, q* was set to 1/(2*N*), and Equation (3a) with binomial sampling was applied for further generations. The same procedure was repeated after the loss of A_2_, allowing new mutations to enter until fixation, loss, or termination.

Simulations were run under varying demographic and selection models, aiming to establish conditions for detecting purging and to determine the time required for its effects to manifest. We mostly considered two different population sizes after a bottleneck, *N* = 25 or 50. The models of selection involved mutations with fixed effects, with selection coefficients ranging from *s* = 0.1 to lethal (*s* = 1). Scenarios included either fully (*h* = 0) or partially (*h* = 0.1 or 0.2) recessive mutations. Nearly-neutral cases (*s* = 0.01 or 0.005) with the same dominance coefficients were also evaluated. Mutation rates (*u*) were calibrated to yield an equilibrium *B* ≈ 4.5 on the basis of the genomic model used in multi-locus simulations (see below), and the resulting values of *u* were used here; these are shown in Table S1 of Appendix S3.

### Population genetic statistics for the single-locus simulations

For each generation of a set of replicates, we calculated the mean frequency of deleterious alleles across all loci,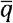; the mean genetic load,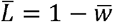; the mean inbreeding load,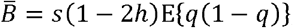; and the fixation index,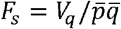. The population statistics 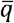, 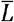 and 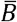 were then expressed relative to their theoretical values for the ancestral population, which are described in the subsection *Population genetic statistics for the ancestral population*; as mentioned above, the subscript *r* is used to denote these relative values. Since these statistics are highly stochastic, a very large number of replicate populations were used to obtain precise estimates of their expected values. Thus, the simulated trajectories represent expectations over many independent evolutionary realizations rather than the trajectory for any single population. To prevent overflow issues, the simulations were usually divided into 25 (*N* = 25) or 15 (*N* = 50) repeat sets of 1 billion replicates, and overall mean statistics were calculated across all repeats and replicates. The trajectories of these expected population genetic parameters over generations were then compared with the expectations derived from the various approximations.

### The multi-locus case

For comparison with single-locus results, and to explore the dynamics expected in genomes with linked sites, we implemented multi-locus simulations in SLiM 4.0.1 (Haller and Messer 2023). In addition to the fixed-effect selection models used in the single-locus analyses, simulations were also performed under an empirically motivated distribution of fitness effects (DFE). Configuration files were adapted from López-Cortegano (2022) and Kyriazis et al. (2023b) (see Data Availability).

Briefly, an ancestral population (*N*_0_ = 1,000) was evolved for 5,000 generations to reach mutation-selection-drift equilibrium, and was then subdivided into smaller populations (*N* = 25 or 50), each run for 100 additional generations. A genome model inspired by mammals (Keightley 2012) was assumed, with 22 chromosomes with map lengths of 100 centiMorgans, each containing 1,000 genes with 1,000 selected sites per gene (i.e., 22 million selected sites per genome). Recombination was assumed to occur only between genes, with no recombination within genes, with a recombination rate *r* = 10^−3^ between adjacent genes and free recombination (*r* = 0.5) between chromosomes. All simulations assumed random mating, and followed the same demographic models as described above for single-locus simulations. Each parameter set was replicated 1,000 times (10 ancestral populations × 100 subpopulations).

The per-generation value of the inbreeding load was calculated as the sum of *B* = *pq*(1 − 2*h*)*s* across loci. Because a large inbreeding load is needed to produce both a substantial fitness decline and purging in a small population, the equilibrium inbreeding load was standardized to *B* ≈ 4.5 in all selection models, in order to facilitate comparisons among them. This value results from the combined effects of the 22 million sites subject to mutation, selection and drift in the simulated genome, and is consistent with previous field and experimental estimates of the inbreeding load (O’Grady et al. 2006; López-Cortegano et al. 2016), which we attribute to deleterious mutations for the purpose of modeling.

Two selection models were considered. In the first, mutations had fixed selection and dominance coefficients corresponding to the single-locus models described above. In the second, mutation effects followed a DFE that was inspired by population genomics inferences (Kyriazis et al. 2023b; Wade et al. 2023). Because this DFE was intended to represent a more realistic biological scenario, relevant to a natural population, these simulations used larger population sizes (*N*_*0*_ = 25,000 and *N* = 1000). Under this model, 0.3% of mutations are fully recessive lethals; the remainder were drawn from a gamma distribution with mean *s* = 1.315 × 10^−2^ and shape parameter 0.186. Following the nomenclature in the original script, semi-lethal (*s* > 0.1, *h* = 0), strongly deleterious (0.01 < *s* ≤ 0.1, *h* = 0.05), moderately deleterious (0.001 < *s* ≤ 0.01, *h* = 0.2) and weakly deleterious (*s* ≤ 0.001, *h* = 0.45) mutations comprised 2.6%, 23.5%, 24.6% and 49.0% of all mutations, respectively.

For the fixed-effects models, the per-nucleotide mutation rate (*u*) was calibrated for each selection model to produce the target equilibrium inbreeding load *B* ≈ 4.5. Starting from the ancestral population (*N*_0_ = 1,000), ten replicate simulations were run for each candidate value of *u*, and the mean inbreeding load over the final ten generations was estimated. Mutation rates were adjusted iteratively to approach the target value as closely as possible, with a target precision of three decimal places. The resulting mutation rates ranged from 2.5 × 10^−9^ to 7.6 × 10^−8^ depending on the selection model (Table S1). These mutation rates were then used in the corresponding single-locus simulations, although the relative values of the mean allele frequency, genetic load and inbreeding load are effectively independent of the exact mutation rate provided that *q\** ≪ 1. For the variable DFE model, the mutation rate was calibrated using the same procedure, yielding *u* = 2.217 × 10^−8^.

Multiplicative fitnesses across loci were assumed, so that the fitness of an individual with *i* heterozygous loci and *j* loci homozygous for A_2_ is (1 − *hs*)^*I*^ (1 − *s*)^*j*^. Under this assumption, locus-specific contributions are additive on the logarithmic scale. Thus, for *l* independent loci with the same selection parameters and with multiplicative fitness effects across loci, the expected value of the natural logarithm of mean fitness is simply 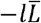 (Haldane 1937). Accordingly, the relative genetic load was calculated as 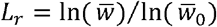, where 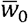 was estimated as the mean fitness of the base population averaged over the last 1,000 generations of the simulations, representing mutation-selection-drift.

## Results

The accuracy of the analytical approximations for strong selection described above, and of the linear differential operator (LO) method described in section 1 of the Appendix, was assessed by comparing them with single-locus simulation results. Table 1 provides an overview of the various methods used for these approximations.

### Comparisons of single-locus simulations and analytical results: strong selection (Methods 1a, 1b, 2a and 2b)

Here, we focus on the short-term (t ≤ *t*_*c*_)consequences of genetic drift and selection on bottlenecked populations, examining their effects on three key population genetic statistics: the mean allele frequency 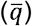, mean genetic load 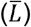 and mean inbreeding load 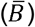, all expressed relative to their equilibrium values (as indicated by the subscript *r*), as well as the fixation index at selected sites ( *F*_*s*_). For all of these parameters, the simulations were in reasonably good agreement with expectations during the generations leading up to the values of *t*_*c*_ predicted by Equation (8), but the approximations often deviate substantially from the simulation results for later generations, especially with *h* = 0.2 see Figures 2 and 3 and Figure S2 of Appendix S4.

**Figure 3.**
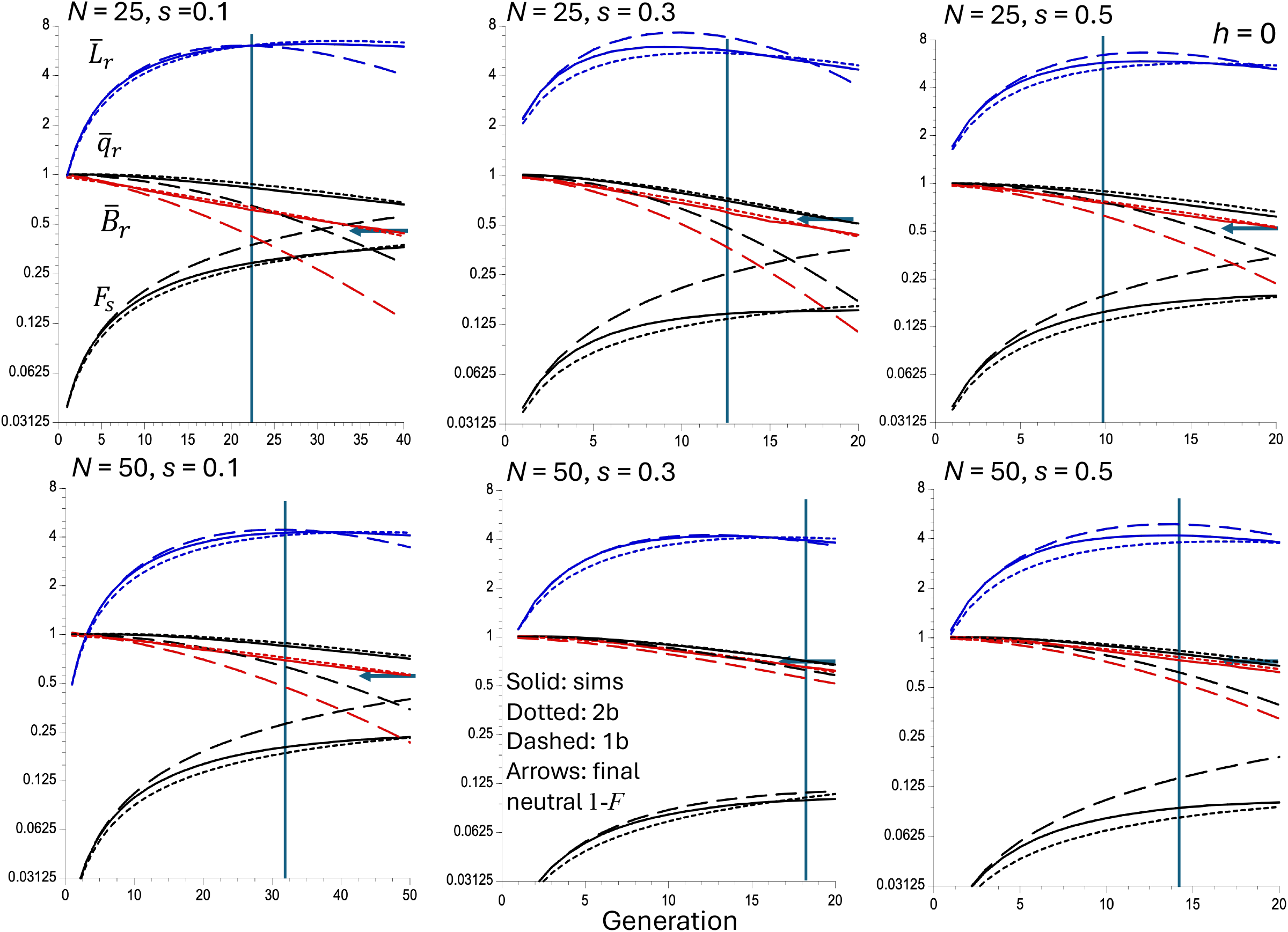
Results of single locus simulations and approximations with *h* = 0 (completely recessive mutations) and strong selection. The Y axis is on a log_2_ scale, because of the large increase in the genetic load in early generations. The solid curves (in order from top to bottom) in each panel show the means of the following statistics over all replicate simulations: the mean genetic load relative to its ancestral equilibrium value 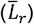, the mean allele frequency relative to its ancestral equilibrium value 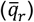, the mean inbreeding depression relative to its ancestral equilibrium value 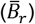 and the mean fixation index at the selected sites (F_s_). The dotted curves are the predictions based on the linear operator Method 2b, and the dashed curves use the approximations given by the trajectories for the mean allele frequency described by Method 1b and the approximation for the relative value of the inbreeding load using Equation (S9a). The arrows display the final value of one minus the neutral fixation index,1 – *F*, which corresponds to 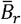 if there is no effect of selection on allele frequencies. The vertical lines indicate the values of the approximate expected time for purging to be manifested (t_c_), obtained from Equation (8) with *h* = 0. See the *Methods of Analysis* section for details of the simulation method.

Equations (4) for 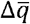 and (8) for c predict a recovery of mean fitness after an initial period of decline associated with the increase in *F*_*s*_, provided that *h* < ⅓. This reflects the fact that the genetic load experiences forces acting in opposite directions – an increase in the fixation index tends to increase 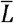(i.e., it reduces the expected population mean fitness), whereas a decrease in 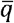 tends to reduce 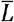. Initially, the first effect outweighs the second, but the reduction in 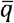 can eventually become sufficiently large to outweigh the effect of an increase in *F*_*s*_, especially as this increase is opposed by selection. In line with these expectations, with *h* < ⅓ the simulations and approximations exhibit a transient increase in 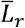 after the bottleneck, which attains its highest value at times that are at most a few tens of generations after the bottleneck. The simulation and analytic results with other parameter sets confirm the expectation that a decline in 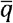 and a recovery of mean fitness are not observed if *h* ≥ ⅓.

The LO predictions (Method 2) in Figures 2 and 3 generally behave better than the more approximate predictions (Method 1) when selection is relatively weak, especially with *h* = 0, but not necessarily with stronger selection and *h* = 0.2. The predictions for *t*_*c*_ from Equation (8) are quite accurate with *h* = 0 (Figure 3), but tend to considerably overestimate *t*_*c*_ compared with the simulation results with *h* = 0.2, especially when *Ns* is large (see Figure 2). Indeed, the simulation results in Figure 2 for *s* = 0.5 and *h* = 0.2 show that 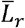 starts to decline immediately after the bottleneck, whereas Methods 1a and 2a predict a decline after approximately 4 generations with *N* = 25 and 50; the corresponding predicted *t*_*c*_ values are 6.31 and 6.81. These inaccuracies probably reflect the fact that the mean fitness denominator in Equation (3a) was not used in the approximate equations, as well as the use of the neutral value of *F* in Equation (8), both of which are especially dubious for large *h* and *s*.

A point of some interest is that, when *h* = 0, Figure 3 shows that the expected load in the first few generations when *s* = 0.1 and *N* = 50 is considerably *smaller* than the load for the initial population, which is approximately equal to *u* (Nei 1968). This effect follows from the fact that the variance in *q* among populations where A_2_ has survived the bottleneck one generation after the bottleneck is approximately *F*_2_(1/2*N*) = (2/2*N*)(1/2*N*) = 2/(2*N*)^2^. The net expected genetic load E{L} = E{q^2^}*s*, in that generation is thus given by:

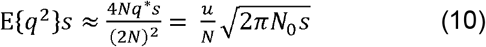

Since the ancestral load is equal to *u*,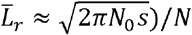. With *s* = 0.1, *N* = 50, *N*_0_ = 1000, this is equal to 0.501, which is close to what is seen in Figure 3. For a small time *t* after the bottleneck, where *Ft* ≈ *t /*(*2N*) this formula is multiplied by *t* /2, so that the genetic load is expected to increase approximately linearly in the early generations. For the other parameter sets in the figure, 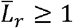 in the initial generations, as predicted by this formula.

In all cases shown here (where *h* < ⅓), both 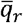 and 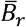 consistently decline over time, as expected from the approximate equations used in Method 1. In relatively large populations (*N* = 50 versus *N* = 25) or with strong selection, 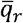 and 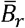 decrease at similar rates, indicating that the reduction in 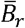 is mainly driven by the decline in 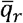. In contrast, smaller populations or those with more weakly selected mutations show a faster decline in 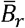 than in 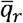, due to loss of variability caused by drift. These results highlight the fact that the increase in variance in allele frequencies (with its associated loss of variability, as reflected in the increase in *F*_*s*_), is only part of the contribution to the decline in 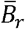. The fact that selection drives down the expected frequency of deleterious alleles also contributes to this decline.

An important point is that the rate of decline in 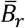 can either be *slower* than expected when using the neutral inbreeding coefficient in Equation (2), or *faster* (with *h* = 0 and the larger *s* values). This result reflects differences in the extent to which selection reduces the rate of increase in *F*_*s*_. In all cases, as expected intuitively, *F*_*s*_ increases over time more rapidly when selection is less effective, i.e., with small *hγ*, where *γ* = 2*Ns*. For *h* > 0, both the Method 2a (LO) predictions for *F*_*s*_ and Equation (S4) closely match the simulation results for the parameter sets considered here, and show an approach to the asymptotic value of *F*_*s*_ given by Equation (S5). For example, bottlenecked populations of size *N* = 50 with *s* = 0.5 and *h* = 0.2 become close to the approximate asymptotic value of 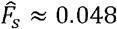 within just ten generations (see Figure 2). For recessive mutations, the Method 2b predictions for *F*_*s*_ were very accurate (see Figure 3), but the neutral prediction tended to greatly overestimate *F*_*s*_. For this reason, the Method 1b approximation for 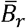 greatly underestimated the simulation values.

For purely recessive mutations (*h* = 0), the approximate value of 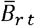 at generation *t*_*c*_ turns out to be nearly independent of *s* and *N*_*e*_. Substituting *t*_*c*_ from Equation (S13c) into Equation (S9b) yields the following result if *T*_*c*_ is sufficiently small that *F*_*s*_ is negligible:

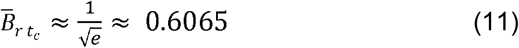

This expression implies that the inbreeding load due to fully recessive deleterious mutations is reduced by roughly 40% of its initial value after *t*_*c*_ generations, regardless of the population size and selection coefficient. This result should, however, be treated with caution, as the simulations and the more exact results with the LO method show that the approximations used here often overestimate the decline in inbreeding load with time. As can be seen in Figure 3, this value is closest to the simulation/LO results for the smaller values of *s* and *N*.

### Comparisons of single-locus simulations and analytical results: weak selection (Methods 2c and 2d)

We now consider the results for cases when selection is so weak that *Ns* for the bottlenecked population is of the order of 1 or less, e.g. for populations with *N* ≤ 10, when *s* is ≤ 0.1 (as pointed out at the end of this subsection, these conditions are in fact too stringent). The single-locus simulation results for these cases can be compared with both the predictions of the more exact weak selection version of the LO method (Method 2c), and with the more extreme approximation that the distribution of *q* is close to the neutral distribution, so that 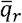 remains close to one and the fixation index increases according to the neutral recursion relation (Method 2d: see section 2 of the Appendix). Intuitively, this approximation yields the expectation that the mean load always increases as *F* increases if *h* < ½, and that 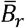 is approximately equal to 1 – *F* (see Equations A21 and A22).

Figure 4 displays the results for the case of *s* = 0.01 and 0.005, and *N* = 25 and 50, with *h* = 0.2. Methods 2a and 2c give extremely good fits to the relevant simulation results; for all but *s* = 0.005 and *N* = 50, they are nearly indistinguishable. The fit is less good for the Method 2d, but still quite close. In all cases, there is no sign of purging at the level of the mean genetic load; instead, it increases monotonically and is close to the value predicted by the neutral approximation of Equation (A21), which reflects the effects of drift alone. 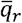 tends to decrease slightly at first and then to increase to slightly above one, reflecting the input of new mutations, but in every case it stays close to one over the time courses used here. Even with this relatively weak efficacy of selection, however, as time increases there are increasing discrepancies between Method 2d and the realised values of the means of the genetic and inbreeding loads, as well as the fixation index, especially for the larger values of *Ns*.

**Figure 4.**
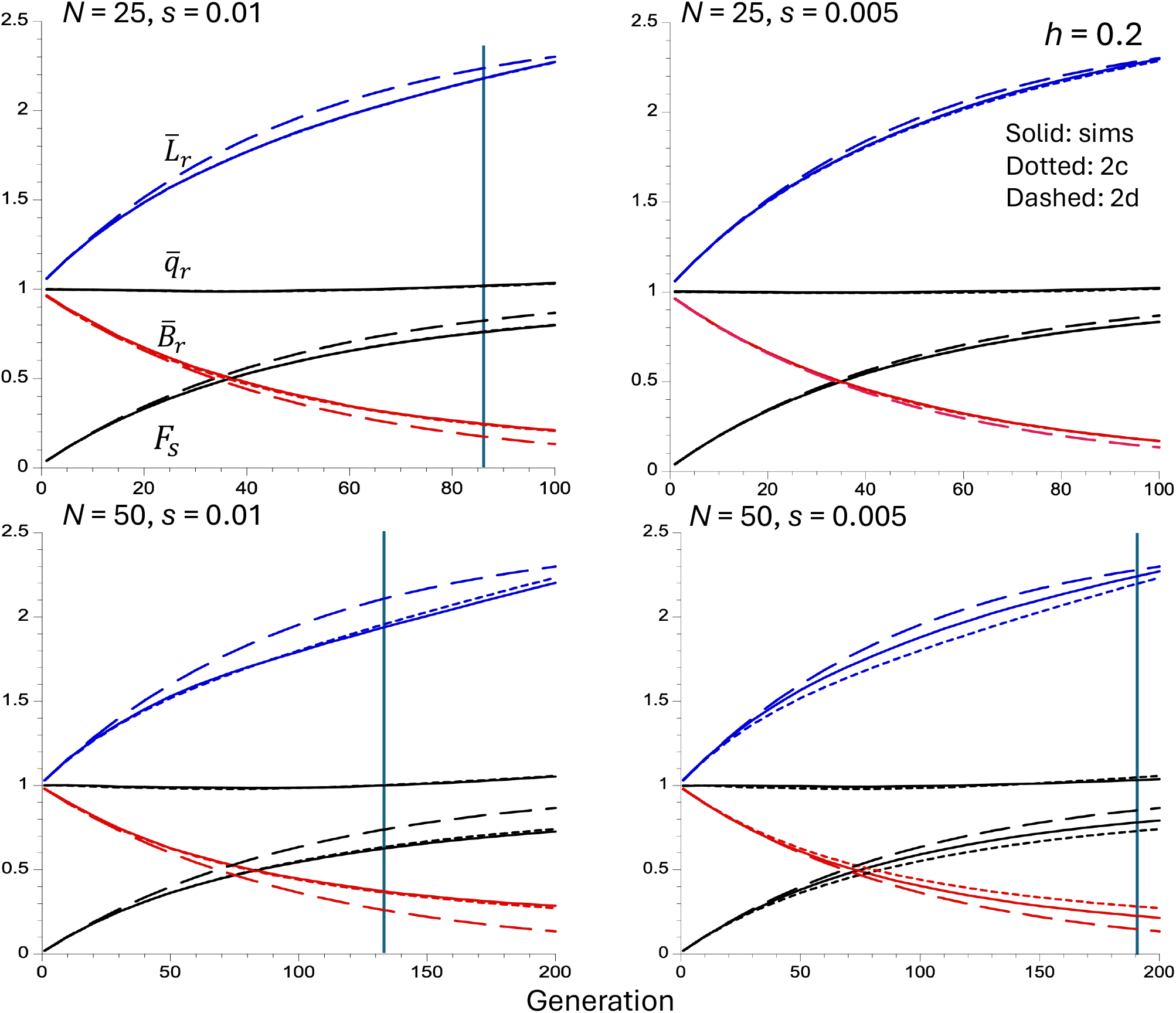
Results of single locus simulations and approximations with *h* = 0.2 and weak selection (*s* = 0.01 and *s* = 0.005 for the left-hand and right-hand panels, respectively). The solid curves (in order from top to bottom) in each panel show the means of the following statistics over all simulations: the mean load relative to its ancestral equilibrium value 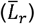, the mean allele frequency relative to its ancestral equilibrium value 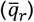, the mean inbreeding depression relative to its ancestral equilibrium value 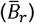 and the fixation index (*F*_*s*_). The dotted curves are the predictions based on the linear operator Method 2c, and the dashed curves are approximations obtained using Method 2d and the approximation for the relative value of the inbreeding load 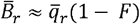. The vertical lines indicate the values of the approximate expected time for purging to be manifested when selection is strong (t_c_), obtained from Equation (8) (t_c_ for *s* = 0.005, *N* = 25 is 142, which is off the scale). See the *Methods of Analysis* section for details of the simulation method.

A point of interest is that the trajectories of the population statistics for *s* = 0.01 and *N* = 25 are close to those for *s* = 0.005 and *N* = 50, if two generations in the latter case are taken to correspond to one generation of the former. This corresponds to what is expected from diffusion equation theory, which predicts that 2*N*_*e*_*s* determines the rate of change of the probability distribution of allele frequencies if time is measured in units of 2*N*_*e*_ generations (Robertson 1960). There are, however, issues with the different initial allele frequencies, conditioned on survival into the bottlenecked population, which complicate this interpretation; these are considered in the Discussion.

It seems plausible that the weak selection LO approximation should perform well if h*γ* << 1 (when h > 0) or *γ* << 1 (when h = 0), where *γ* is the scaled selection coefficient, since these correspond to conditions when drift is strong or weak relative to selection as far as the fate of new mutations is concerned (Crow and Kimura 1970). Conversely, the strong selection LO approximation should be accurate if h*γ* >> 1 (when h > 0) or if *γ* >> 1 (when h = 0). However, it is unclear exactly where the lines should be drawn. This question was examined by running single locus simulations with *N* = 25 for a set of different s and h values, and comparing them with the results of the corresponding weak and strong LO predictions (see Appendix S5).

Figures S3 and S4 show results for h = 0.1 and h = 0.2, for four different s values, chosen such that the four panels in each figure correspond to h*γ* = 0.25, 0.5, 0.75 and 1, respectively. It is evident that the weak selection approximation performs best with h*γ* = 0.25 but worst with h*γ* = 1, whereas the opposite is true of the strong selection approximation, as would be expected. Both LO Methods 2a and 2c work quite well with *h* = 0.2 for all the values of h*γ* covered except with respect to 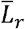, although 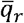 tends to be slightly above one in later generations, whereas the simulated values are less than one. With *h* = 0.1, the weak selection Method 2c performs poorly for 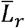 and (to a lesser extent for)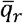, except for h*γ* = 0.25, whereas the strong selection approximation is quite accurate for all values of h*γ*

Figure S5 shows results for *h* = 0 with *γ* = 1.25, 2.5, 3.75 and 5. In this case, the LO Method 2c performs well except for *γ* = 5, surprisingly, the strong selection approximation performs well for all the *γ* values that were investigated.

These results suggests that, for partially recessive deleterious mutations, it is safe to use the weak selection LO approximation (Method 2c) for h*γ* ≤ 0.25, and the strong selection LO approximation (Method 2a) for h*γ* ≥ 1; for intermediate values, Method 2a is on the whole to be preferred, although it gives inaccuracies with respect to 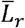 with the larger of the two *h* values. For completely recessive mutations (*h* = 0), the LO Method 2c is accurate for *γ* ≤ 1.25; the strong selection LO Method 2b can be used for *γ* > 1.25.

### Multi-locus simulation results

To examine the extent to which linkage modifies the population genetic dynamics predicted by the single-locus models, multi-locus simulations with a set of 22 million loci distributed over 22 independent chromosomes, each with 1 million selected loci and a map length of 100 centiMorgans were, carried out (see the subsection *The multi-locus case* above for details of the genome model and selection regimes). The results for simulations assuming the empirically motivated DFE are shown in Figure 5, and simulations assuming fixed selection coefficients are shown in Figures S6-S8 of Appendix S6.

**Figure 5.**
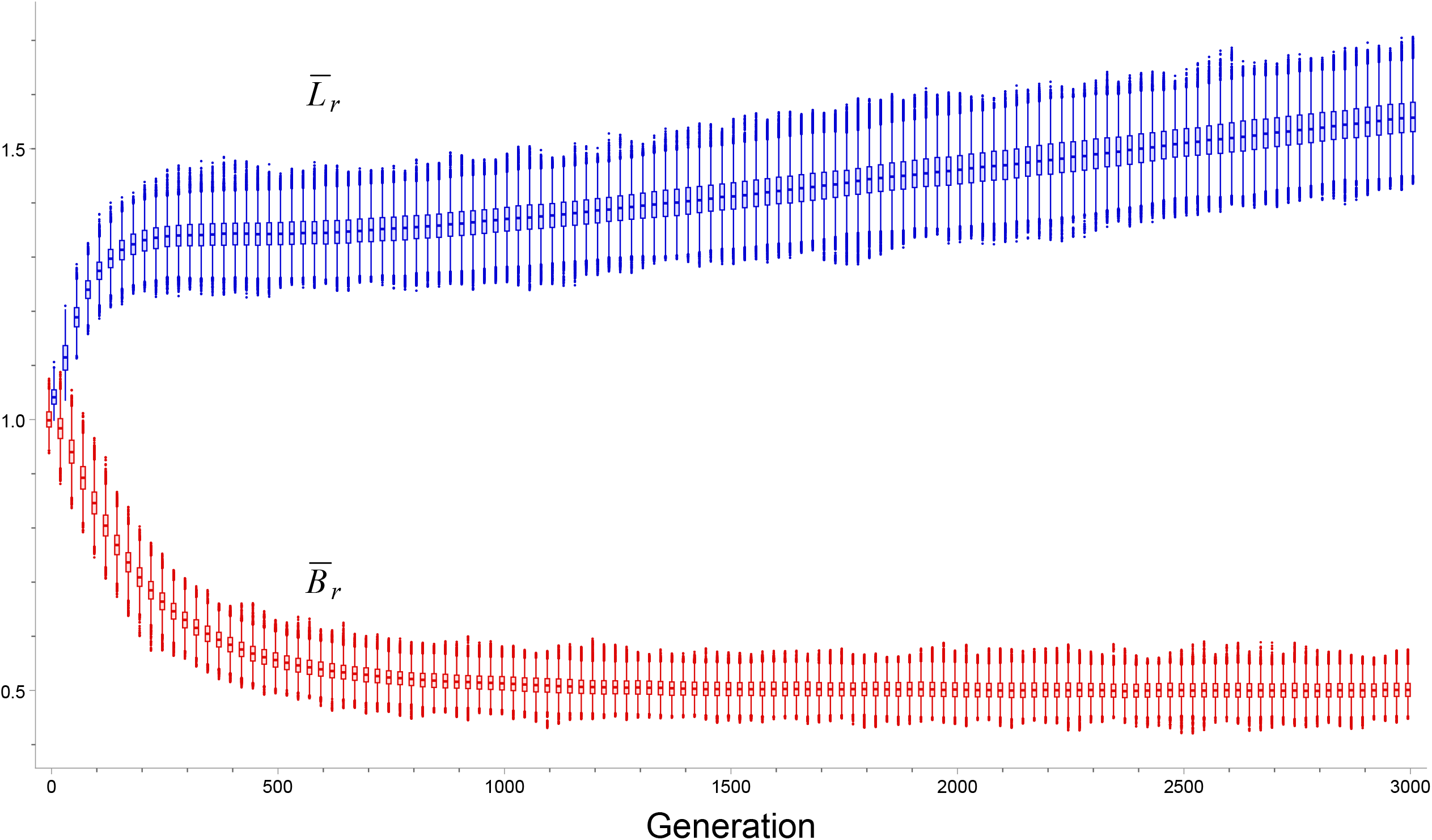
Results of multi-locus simulations with a variable DFE (*N*_*0*_ = 25,000, *N* = 1000). Boxplots show the distributions across simulation replicates of the mean load and inbreeding load relative to their ancestral equilibrium values. Boxes indicate the median and interquartile range (IQR), whiskers the range 1.5 × IQR, and points beyond the whiskers indicate outliers. The relative genetic load 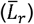 is shown in blue, and the relative inbreeding load 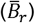 in red. For visualization purposes, generations are rounded to the nearest 25 generations.

The DFE used in the simulations captures a broad spectrum of deleterious mutations of the type expected for a natural population that has gone through a moderate population size bottleneck (Kyriazis et al. 2023b). It is dominated by weakly deleterious mutations with *γ* values close to 1 and *h* values that were much greater than zero but less than 0.5, although a small fraction of mutations were very strongly selected and had *h* values of zero or close to zero. The mean of the population fitness over all replicates 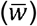 declined from an equilibrium value of 0.59 in the ancestral population (*N*_*0*_ = 25,000) to 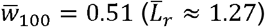 after 100 generations. The rate of decline in 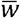 subsequently slowed substantially after approximately 200 generations, reaching a minimum rate at 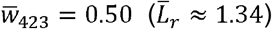, suggesting that purging of the most strongly deleterious mutations involved in the simulations (*s* > 0.01 and *h* ≤ 0.05; corresponding to *t*_*c*_ < 500) contributed to the attenuation of the rate of decline in 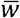 After an almost undetectable transient recovery, however, 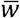 resumed a nearly linear, slow decline for the remaining time period 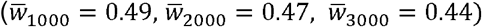, corresponding to 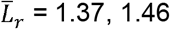 and 1.56, respectively. Thus, although strongly deleterious mutations can contribute to short-term purging, in this case their removal was insufficient to compensate for the continued accumulation and persistence of weakly deleterious variants. Importantly, the box plots show that there is relatively small variation in both 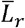 and 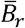 among replicate simulations, suggesting that their overall means provide a reasonably good guide to what would be observed in a single population.

To facilitate direct comparisons with the single-locus simulations and analytical models, additional simulations assuming fixed mutational effects were also conducted. However, it should be noted that the multi-locus design used here, with millions of sites and fixed selection coefficients (e.g., *s* = 0.1, 0.5, etc), strongly exaggerates the potential effects of linkage, given the evidence from population genomics studies that most deleterious mutations have selection coefficients of the order of 0.01 or less (Kim et al. 2017). These simulations therefore represent an extreme test of the robustness of the single-locus approximations, and were mainly aimed to test the accuracy of the conditions for detecting purging and its expected time span as given by Equation (8). It is thus not surprising that both 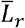 and 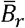 tended to be somewhat higher in the multi-locus simulations than in the single-locus simulations, especially for the smaller population size (Figures S6 and S7), with these differences becoming more pronounced over time. This pattern might be expected if Hill-Robertson effects induce pseudo-overdominant associations among loci segregating for recessive or partially recessive deleterious mutations, which persist and accumulate over the long-term (see the Discussion). Despite these long-term differences, the trajectories of 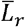 in the short-term (*t* < *t*_*c*_) were generally similar under the single and multi-locus models, especially for *s* = 0.1 and *h* = 0.2, with only small differences that sometimes changed sign over time.

A fitness recovery was not always observed in these simulations, suggesting that strong linkage could substantially reduce the efficiency of purging relative to single-locus expectations, and effectively increase the *Nhs* ≥ 1 threshold described above almost ten-fold, due to extreme effects caused by linkage disequilibrium. In cases where mean fitness recovered after an initial period of inbreeding depression, the observed timescale of recovery generally agreed with the expectation derived from Equation (8). The largest differences between the simulated and analytical estimates of *t*_*c*_ were observed for partially recessive mutations (*h* = 0.2). In these cases, Equation (8) consistently underestimated *t*_*c*_, indicating that purging required somewhat more generations to become manifest in the multi-locus than predicted analytically. However, this effect was modest, typically amounting to fewer than ten generations

## Discussion

### Some general considerations

We have developed two new approaches to modeling the behaviour of rare, deleterious, autosomal mutations in a bottlenecked population. These were used to predict the approximate time courses of the expectations of the frequencies of deleterious alleles, the genetic load, and the inbreeding load in a small population founded from a large population that was close to mutation-selection-drift equilibrium. Our results show that these properties of a bottlenecked population reflect three familiar consequences of a reduction in population size to *N* from a much larger ancestral size, *N*_0_. The first is the loss of rare deleterious alleles, which occurs with a probability of approximately *P* = 1 – 2*Nq*\*, where *q*\* is the equilibrium expected frequency of mutant alleles. The second, with approximate probability 2*Nq*\*, is the presence of the surviving mutant alleles at an elevated expected conditional frequency of 1/(2*N*). The third factor is the variance in allele frequencies across different loci induced by the reduced population size, which increases the expected frequencies of homozygotes over Hardy-Weinberg expectations.

The last two factors should in principle should intensify the effectiveness of selection against rare recessive or partially recessive alleles, although emphasis is usually placed on the third factor (e.g. Glémin 2003). Our results show that a decrease in the short-term expected net frequencies of rare deleterious alleles following a bottleneck requires both *h* <⅓ and sufficiently strong selection that the final stationary state under mutation, selection and drift has a low mean frequency of deleterious mutations (see Kimura et al. 1963; Lande 1998; Charlesworth 2018). The condition on *h* has not previously been explicitly described in relation to a bottleneck, as far as we are aware, although it follows from Equation (11) of Whitlock (2002) for the change in expected allele frequency in a subdivided population. In their study of bottlenecks, however, Kirkpatrick and Jarne (2000, p.158) stated that *h* < 0.3 was required for purging, on the basis of numerical results.

However, with *γh* < 0.25 for partially recessive mutations, or *γ* < 1.25 for completely recessive mutations (where *γ* = 2*Ns*), genetic drift tends to dominate over selection and the expected mean frequency of deleterious alleles remains approximately constant in the short term, but increases slowly under mutation pressure. The mean genetic load increases, and the mean inbreeding load decreases, at rates that are close to what is expected on the basis of a neutral increase in the fixation index (see Figures 4, S3 and S4). With *h* > 0 and *γh* between 0.25 and 1, the behavior of the population statistics depends on *h*, with smaller values allowing a decrease in mean allele frequencies and genetic load even when *γh* < 1 (Figures S3 and S5).

### Conditions for purging of deleterious mutations

A more exact condition for purging at the level of allele frequencies with strong selection (*γh* > 1, when *h* > 0) is provided by Equation (A12) for the final equilibrium value of the expected allele frequency, which uses the linear operator Method 2a (strong selection). This expression implies that this frequency is less than the value for the ancestral population if and only if:

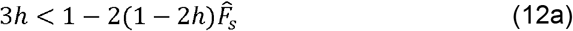

where 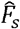 is given by Equations (A9) and (A10), or more approximately by Equation (A11). A less accurate expression for 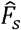 is:

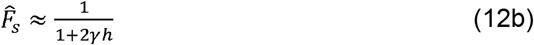

This expression assumes that 2 *γ h* = 4*Nhs* ≫ 1, implying that 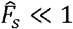. Given that 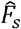 is a decreasing function of *γ* and that (1 − 2*h*) > 0 when *h* <, the maximum value of *h* that satisfies Equation (12a) must be a monotonically increasing function of *γ*. Appendix S7 analyses the solution to the quadratic equation for *h* given by combining Equations (12a) and (12b). This solution is meaningful only if *γ* > 4.95, corresponding to *h* < 0.184. Above this threshold, the larger *γ*, the larger the corresponding *h*, reaching an asymptote of *h* = ⅓ very quickly.

This analysis is consistent with the fact that examination of the exact numerical results for the equilibrium distribution of allele frequencies with population size *N* shows that a reduction in the final mean frequency of deleterious mutations below their ancestral values requires *γ* >>1 for partially recessive mutations, even with *h* < ⅓. For example, with *N* = 50, *s* = 0.1 and *h* = 0.2 (*γ* = 10) (the values used in the left-hand panels of Figure 2), numerical solutions of Equation (S7) of Charlesworth (2018) yield equilibrium values of 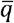, 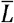 and 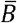 relative to their large population values of 39.6, 97.9, and 0.700, respectively, assuming that reverse mutations from A_2_ to A_1_ occur at one-third of the rate of forward mutations. Inclusion of reverse mutations is essential for the system to reach a final equilibrium under drift, mutation and selection; in their absence, there will simply be an irreversible accumulation of deleterious mutations (Wright 1931). This contrasts with values of 0.861, 1.04, and 0.741 from the approximate Equations (A11) and (A12). But with *N* = 100, the final equilibrium relative values of 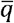, 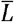 and 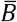 are 0.840, 1.00 and 0.760, respectively, compared with the approximate predictions of 0.887, 1.00 and 0.817, so that the strong selection LO approximation (Method 2a) then gives a reasonably accurate description of the final state (reverse mutations have little effect on the outcome when the expected frequency of A_2_ is low).

### Conditions for the genetic load to be reduced below its initial value after a bottleneck

With strongly selected and highly (but not completely) recessive mutations, 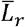 can become substantially smaller than one. For example, with *s* = 0.5, *h* = 0.2 and *N* = 50, the simulations show that 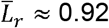 after 15 generations (see Figure 2, bottom right), which is close to the final equilibrium value of 0.88 estimated from long-term simulations, while the Method 2a approximation gives 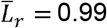, reflecting a predicted 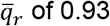 compared with the simulation value of 0.82. The reduction in genetic load below the initial value is even more marked with *h* = 0.1 (see Figure S2). In this case, with *s* = 0.3 or 0.5 the relative expected inbreeding load 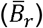 during the early generations after the bottleneck is usually (but not always) less than that expected on the basis of the increased homozygosity of neutral variants, which is equal to the neutral 1 – *F*, reflecting the effect of selection in removing deleterious alleles. For example, with *s* = 0.5, *h* = 0.2 and *N* = 50, 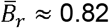 at generation 15, whereas neutral 1 – *F* = 0.86.

A condition for the final equilibrium genetic load to be less than that for the ancestral population when *h* > 0 can be found using the results in section 1 of the Appendix for the linear operator Method 2a, in a similar way to that used above for the mean equilibrium deleterious allele frequency. As shown in Appendix S7 of the Supplementary Material, to the order of the approximations used to derive the equilibrium conditions we have 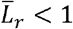 if and only if:

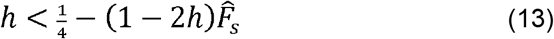

For 2 *γ h ≫*, so that 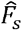 is close to zero, this is equivalent to 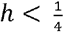, a stronger condition than that for 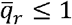, reflecting the effect of increased homozygosity in increasing the load when *N* is small. When Equation (12b) is used as an approximation for 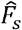, a similar analysis to that used above for 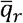 shows that the condition on Equation (13) implies that *h* ≤ 0.138 and *γ* > 19.8 are required for 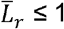; the critical value of *h* increases monotonically towards ¼ as *γ* increases (see Appendix S7).

### Relations to other recent theoretical studies

It is useful to relate our results to other recent approaches to modeling the effects of bottlenecks. An influential method for analysing the effects of bottlenecks with strong selection was proposed by García-Dorado (2012), hereafter GD, using the concept of the “purged inbreeding coefficient” (*g*) as a heuristic tool (reviewed by Hedrick and García-Dorado 2016). In our notation, 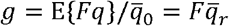, where F is the neutral fixation index. To use this parameter, the equation for expected allele frequency (our Equation 3c) is approximated by the expression 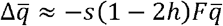 by ignoring the term in 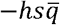, which leads to a simple recursion relation for *g* in terms of *N* and *F* (Equation 7b of GD). In the absence of new mutations, the expected inbreeding load at a given time is given approximately by 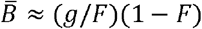 (Equation 9 of GD), and the difference between the population mean fitness for a single locus and its initial value is equal to 0, where 0 is the initial value of the inbreeding load (Equation 3 of GD). Additional terms can be added to include the effects of new mutations, and to calculate the asymptotic values of the mean fitness and inbreeding load (Equations 11-14 of GD).

This approach effectively collapses the effects of selection and dominance into a single parameter 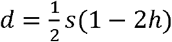, whereas the present approach treats *s* and *h* explicitly and does not neglect a potentially important term in the basic selection equation. While the GD method appears to yield reasonably accurate short-term predictions of mean fitness (e.g.,López-Cortegano et al. 2016), it does not accurately capture some features of the effects of bottlenecks. For example, no increase in mean fitness over that for the initial population is predicted, contrary to what happens with strong selection when 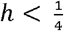 and *γ* is large (see above). The method is also not capable of dealing accurately with the behavior of bottlenecked populations when *γ* is sufficiently small that drift has a major effect on the final outcome, as in the case of *s* = 0.1 and *h* = 0.2 in Figure 2 and the parameters used in Figure 4.

Balick et al. (2015) presented a theoretical analysis of the effects of a bottleneck on variants at sites subject to purifying selection that is closer in spirit to ours than GD, using the forward diffusion equation, supplemented by single-locus simulations. They assumed that the population remains at a reduced size equivalent to our *N* for *T*_*B*_ generations, and then re-expands instantaneously back to the initial size *N*_0_. They derived approximations for the dynamics of the population statistics over the time period following the recovery, assuming that the effect of the first generation of a bottleneck, with the addition of a term reflecting the contribution of new mutations, predicted the moments of the distribution of allele frequencies at the end of the bottleneck. Their measure of the “relative mutational burden”, *B*_*R*_, is equivalent to the inverse of our 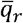.

The results of their model are thus not directly comparable with ours, since we did not model a population re-expansion. However, their qualitative conclusions are similar to ours, in that an increase in *B*_*R*_ over 1 (equivalent to 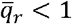) required *h* to be less than a critical value, *h*_*c*_, and was most marked when *h* = 0. They estimated that *h*_*c*_ ≈ 0.25, whereas our analysis suggests that *h*_*c*_ ≈ 0.33 (see above). *B*_*R*_ at a time *t*_*obs*_ after the bottleneck was highest for a given *h* for large values of *s* for small values of *t*_*obs*_, but was smallest for low values of *s* when *t*_*obs*_ was large. The behavior of *B*_*R*_ for small *t*_*obs*_ is similar to that for 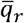 in Figures 2 and 3; its behavior for large *t*_*obs*_ reflects the fact that larger *s* results in a faster approach to mutation-selection equilibrium.

Somewhat similar conclusions were reached by Simons et al. (2014) using computer simulations of several different demographic models, including a simple population bottleneck, but assuming either no dominance or complete recessivity (*h* = 0). Recessivity was required for a reduction in the mean number of deleterious mutations per haploid genome during a bottleneck, and was accompanied by a transient increase in the genetic load, which is proportional to the frequencies of homozygotes in the case of complete recessivity (see our Figure 3).

None of these investigations examined the inbreeding load. Kyriazis and Lohmueller (2024) used multi-locus simulations of nonsynonymous mutations across the human genome to compare the properties of African populations with a transiently bottlenecked out-of-Africa (OOA) population with a wide range of assumptions about selection and dominance coefficients. They found that the genetic load was slightly reduced in the OOA population when deleterious mutations were strongly recessive, consistent with the effects of purging of mutations when a reduced population approaches its long-term equilibrium under strong selection and *h* is small. However, the opposite was the case for weakly or moderately recessive mutations, probably because of the effects of deleterious mutations drifting to fixation when *γ* is small. They found parallel, but somewhat weaker, patterns for their equivalent of 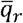, the mean derived allele count per individual. Their theoretical values of the inbreeding load showed no differences between the two populations for weakly or moderately recessive mutations, and a modest reduction for strongly recessive mutations. These relatively minor effects compared with those seen in our Figs 3-5 probably reflect the relatively weak bottleneck that was modelled (*N*_0_ = 1861 for 1120 generations), such that neutral *F* reached only 0.26, as well as a larger final population size for the OOA population compared with the African population. These differences in findings bring out the fact that the theoretical expectations for the effects of bottlenecks are strongly dependent on the details of the demographic model involved, as well as on the selection parameters.

Other theoretical studies of the effects of bottlenecks, with especial reference to OOA populations, include Koch and Novembre (2017), who assumed a purely semidominant selection model and focussed on allele frequency spectra derived from numerical solutions of the forward diffusion equation. Gravel (2016) also used the forward diffusion equation to obtain recursions for the moments of the allele frequency distribution. He examined various measures of the genetic load that differ from Crow’s definition used here (*L*), as well as one that is equivalent to *L*. His numerical results showed that fully recessive or partially recessive (*h* = 0.2 or 0.05) mutations could lead to a reduction in *L* for the OOA population following an initial increase, especially when mutations are highly recessive, consistent with our results.

For modeling the human bottleneck with our approach, and assuming some commonly used parameters in the literature, such as *N* = 1000, *s* = 10^−3^, *h* = 0.2; *hγ* = 2*Nhs* = 0.4, the best approximation is probably Method 2a, which predicts no recovery in mean fitness after a bottleneck for *γ* values of this order (Figure S4). Direct extrapolation of this equation to humans is nevertheless difficult because our model assumes a sustained bottleneck without a subsequent population expansion, and this calculation ignores any contribution from strongly deleterious and highly recessive mutations. Overall, however, the results suggest that genetic purging is unlikely to make a substantial contribution to fitness recovery in our species, but further modeling with a realistic distribution of selection coefficients and dominance coefficients, as well as allowing for possible effects of linkage, is required to reach a definitive conclusion.

Brandvain and Wright (2016) have advocated the use of the ratios of the values of polymorphism statistics for putatively selected versus neutral sites (e.g., nonsynonymous and synonymous sites, respectively) to characterize a potential reduction in the efficacy of selection in a population that has experienced a reduction in size. If π_*sel r*_ and π_*neut r*_ are the ratios of the expected values of the diversities at selected and neutral sites in a population with reduced size to their values in a larger population, the criterion 1 < π_*sel r*_ /π _*neut r*_ is taken to indicate a reduced efficacy of selection. Use of estimates of this diversity ratio, or the corresponding ratio of estimates based on the numbers of segregating sites, as a test statistic is a common practice in population genomics studies – see Table 1 of Brandvain and Wright (2016).

The analysis in Appendix S10 shows, however, that there are difficulties with this approach, somewhat different from those pointed out by Campos et al. (2014). The overall conclusion is that it is easier to satisfy 1 < π_*sel r*_ /π _*neut r*_ than 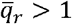. The condition 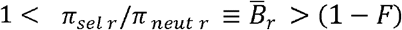 can be met even with purging of deleterious alleles 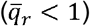, provided that *F*_*S*_ is sufficiently small compared with *F*, Ironically, this condition means that selection is sufficiently strong that the dispersion of allele frequencies at the selected sites is substantially smaller than at neutral sites.

The two left-hand and central panels in Figure 2 exemplify this situation. In contrast, the lower right-hand panel, with strong selection and large *N*, show that is slightly smaller than 1 − *F*, corresponding to1 > π_*sel r*_ /π_*neut r*_, even though than 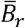 and 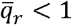 continues to decrease over the time period in question. When selection is very weak, as in Figure 4, 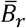 is close to 1 − *F* and 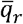 is close to 1, so that there would be no detectable change in the efficacy of selection during the initial stages of the bottleneck on the diversity criterion or the condition 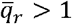, despite a massive reduction in *N*_*e*_*s*. As discussed in the next subsection, the mean genetic load associated with a given *s* and *h* can also start to decrease after a bottleneck when1 < π_*sel r*_ /π _*neut r*_, after an initial increase. As noted by Gravel (2016), clarity about the meaning of terms like “reduced efficacy of selection” is needed when making inferences from population genomics data.

### The expected time to a recovery of mean fitness

Our approximate Equation (8) for *t*_*c*_, the critical time for expected population mean fitness to start recovering, provides a heuristic criterion for distinguishing between the short- and long-term consequences of drift, selection and mutation for the genetic and inbreeding loads. Assuming *h* > 0 and *s* ≫ 0, *t*_*c*_ as given by Equation (8) reaches a limit of (1 – 2*h*)/[2(1 – 3*h*)*hs*] as *N* increases (see Figure S9 of Appendix S8). At this limit, *t*_*c*_ always decreases with increasing *s*; differentiation with respect to *h* and solving the resulting quadratic equation shows that it has a minimum at *h* ≈ 0.21. This reflects a balance between two opposing forces: purging becomes more effective as deleterious alleles are increasingly exposed to selection against heterozygotes (*i*.*e*., with larger *h*; see Equation 3c), but the potential for fitness recovery is progressively reduced as *h* increases (and eventually lost for *h* ≥ ⅓) because of the greater effect on mean fitness of mutations with larger *h* for a given allele frequency.

Because Equation (8) relies on a neutral approximation for *F*, it tends to overestimate *t*_*c*_ with large *h* compared with the single-locus simulations (see Figures 2 and 3) when *Ns* is sufficiently large that a mean fitness recovery occurs. In contrast, Figure S8 shows that it provides a lower bound in the multi-locus simulations with fixed selection coefficients. The reason for this difference is unclear, but may reflect the effects of selective interference in reducing the effectiveness of selection in the multi-locus simulations.

Nevertheless, these biases are modest and do not alter the conclusion that purging occurs over the scale of less than *N* generations, if it occurs at all. Although the asymptotic approximation is not valid for *h* = 0, the order-of-magnitude timescale that it suggests remains a useful expectation for strongly deleterious, highly recessive mutations. These are plausible conditions, given that strong selection is required for purging (see above, and Wang et al. 1999; Glémin 2003), and that the mean value of *h* for minor-effect mutations appears to be approximately 0.25 (Crow 1993; Di and Lohmueller 2024).

Previous approximations for *t*_*c*_ (López-Cortegano 2020) were based on the fraction of the genome exposed to inbreeding in ancestors (*F*_*a*_, Ballou 1997), because heuristic theory suggested that individuals with higher *F*_*a*_ should be more fit than those with equivalent inbreeding but lower *F*_*a*_ due to purging (Boakes et al. 2007). Those approximations yielded 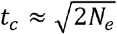 from the generation at which the rate of increase in *F*_*a*_ is maximal. The rationale was that, because *F*_*a*_ rises quadratically over time (López-Cortegano et al. 2018), the opportunity for purging to become detectable should be concentrated within a relatively narrow time window. However, because this approach did not take into account the effects of both *s* and *h*, it was effectively limited to predicting *t*_*c*_ for fully recessive lethals (see Equation (S13c)). In contrast, our model explicitly incorporates these parameters, albeit with some approximations, providing a more precise description of the time required for purging to become manifest under varying genetic architectures (Figures 2, 3 and S8).

### Relations to empirical studies of mean fitness and inbreeding depression

These theoretical results provide a useful reference point, but relating them to the observed trajectories of fitness traits in laboratory experiments on the effects of bottlenecks (mostly involving *Drosophila melanogaster*) is not straightforward. In particular, the extent to which there is evidence for purging from a recovery in mean fitness differs across studies (Latter et al. 1995; Swindell and Bouzat 2006; Ávila et al. 2010; Pekkala et al. 2012; López-Cortegano et al. 2016, Pérez-Pereira et al. 2021). For example, Swindell and Bouzat (2006) found a reduction in inbreeding depression and increased mean fitness for mean offspring production in *D. melanogaster* populations maintained at a small size of *N* = 10 breeding pairs for 19 generations, and Pekkala et al. (2012) found that *D. littoralis* populations maintained at *N* = 10 individuals showed an initial recovery in mean offspring production after five generations, followed by a decline. Populations maintained at *N* = 40 showed a slower decline without an initial recovery. This contrasts with results for competitive fitness over replicate populations of *D. melanogaster* maintained at *N* = 80, which showed an initial decline followed by a recovery (López-Cortegano et al. 2016). However, Latter et al. (1995) observed a decline in net mean fitness measures for replicate *N* ≈ 70 populations of *D. melanogaster* after approximately 160 generations, with no evidence of a recovery.

Regarding inbreeding load, López-Cortegano et al. (2016) found evidence for a considerably faster rate of decline in inbreeding depression in their populations than was expected in the absence of purging. Pérez-Pereira et al. (2021) extended this experiment beyond 100 generations and found that their estimate of inbreeding load was virtually zero. Under a similar rate of inbreeding (Δ*F* ≈ 1%) and timespan, Latter et al. (1995) used a 2^nd^ chromosome balancer and found a large reduction in inbreeding load for detrimentals and lethal chromosomes in their small populations; however, their results did not indicate complete exhaustion of the inbreeding load.

The contrasting outcomes for the effects of experimental bottlenecks on measures of mean fitness suggest that the consequences of bottlenecks depend on factors such as the sizes of the experimental populations, the species and its ancestral population history, the number of generations involved, and the method of estimating fitness. An additional problem is that comparisons of fitness measures across successive generations are likely to be influenced by uncontrolled environmental factors, as well as adaptation to the laboratory environment of the wild-derived flies used in the experiments, e.g. Latter and Mulley (1995).

A major problem in interpreting these experimental results is the fact that the distribution of mutational effects on fitness (the DFE) is dominated by a large majority of mutations subject to relatively weak selection (Eyre-Walker and Keightley 2007; Boyko et al. 2008; Kim et al. 2017, Zeng et al. 2021; Vaughn and Nielsen 2024). Mutations subject to this strength of selection will behave like those modeled in Figure 4 when a population is put through bottlenecks of the sizes used in the laboratory experiments, *i*.*e*., they will behave close to neutrally.

Consistent with the findings from population genomics, studies of the homozygous fitness effects of chromosomes extracted from natural populations of various *Drosophila* species have shown that there is a sharp division into homozygous lethal and sterile chromosomes versus chromosomes with relatively minor reductions in fitness (Simmons and Crow 1977; Crow 1993). In addition, studies of the rates of mutation to lethal and sterility mutations in *Drosophila* (Crow 1993) indicate that these are generally only a small fraction of the total genomic mutation rate, which is now well-characterized from sequencing studies (e.g. Wang et al. 2023). Since the mean autosomal genetic load in large natural populations that is derived from partially recessive deleterious mutations is independent of the strength of selection, and is equal to approximately twice the mutation rate *u* (Haldane 1937), only a small fraction of the genetic load is likely to be caused by these major effect mutations, and their expected contribution to the change in mean fitness following a bottleneck will be relatively minor. Predictions of the effects of a bottleneck of the size used in laboratory experiments therefore requires integration of information on the DFE for minor effect mutations with that concerning lethal and sterile mutations; this will the subject of a subsequent paper. The present qualitative considerations suggest, however, that a recovery in mean fitness is usually likely to be small or non-existent when the whole spectrum of deleterious mutations is taken into account.

However, the inbreeding load *B* when 2*Ns* >> 1 is inversely related to the dominance coefficient, since the equilibrium value of *B* for partially recessive mutation in an infinite population 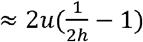 (Charlesworth and Charlesworth 1987), so that highly recessive mutations contribute disproportionately to *B*. In *Drosophila*, homozygous lethal mutations with a mean *h* of approximately 0.02 contribute to about half of *B* for egg-to-adult viability (Simmons and Crow 1977; Crow 1993), despite their small contribution to the genomic mutation rate. Their purging from small populations is thus likely to cause a detectable reduction in *B*, consistent with the observations discussed above.

This conclusion is reinforced by our multi-locus simulations incorporating a DFE informed by population genomic studies (Kyriazis et al. 2023b, Wade et al. 2023) – no fitness recovery was observed after a bottleneck (Figure 5), as expected from an inbreeding load dominated by weakly deleterious mutations. There is, however, a problem with relating single locus results to multi-locus models, since linkage disequilibrium (LD) among the selected sites may cause Hill-Robertson interference (Hill and Robertson 1966), reducing the efficacy of selection and accelerating the loss of variability. In addition, provided that *Ns* is sufficiently small and linkage is sufficiently tight, negative LD among recessive or partially recessive deleterious alleles at different loci could result in pseudo-overdominance, reducing the rate of purging (Latter 1998; Charlesworth 2012; Zhao and Charlesworth 2016, Bersabé et al. 2016; Waller 2021). These effects of LD presumably explain the generally higher genetic and inbreeding loads observed in the multi-locus simulations compared to those using single-locus dynamics (Figures S6 and S7), particularly in the long term. However, the approximation for *t*_*c*_ was reasonably accurate for most of the cases considered in (Figure S8), and the values of 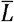 and 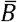 for single and multi-locus simulations agreed quite well before *t*_*c*_ was reached (Figures S6 and S7).

### Purging at the level of the mean number of mutations per genome

Population genomic studies allow estimation of the “burden” of deleterious mutations, defined as the mean number of mutations classed as deleterious or potentially deleterious per haploid genome in individuals sampled from a population (Balick et al 2015). Changes in the burden following a bottleneck result from changes in our 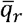, as the burden relative to that for the ancestral population is equal to the product of 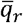 and the number of genomic sites in question. As discussed in the subsection *Conditions for purging of deleterious mutations*, a noticeable reduction in 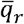 is expected only for mutations that are sufficiently recessive that *h* < ⅓ and that have 2*Nhs* ≫ 1 in the bottlenecked population. We have mainly been concerned with modeling very small laboratory populations; the majority of variants detected in resequencing studies in such experiment will behave like those in Figure 4, *i*.*e*., at most there will be a very small increase in the burden over time as new mutations accumulate.

There is, however, much interest in the population genomics of natural populations that have suffered recent, drastic reductions in numbers, as well as the more minor bottleneck experienced by out-of-Africa humans discussed above (for a recent review, see Robinson et al. 2023). Here, *N* is often of the order of several hundred or several thousand. This raises the question of the relevance of our results for such large *N* values. Diffusion equation theory, which we used in the guise of Method 2, implies that evolutionary change depends primarily on the products of the deterministic parameters *s* and *u* with the variance effective population size (2*N*_*e*_), measuring time in units of 2*N*_*e*_ generations (*i*.*e*., the coalescent timescale) (see Chapters 4 and 5 of Ewens 2004, and Johri et al. 2026).

Rescaling in this way implies that populations differing in absolute size but sharing identical scaled parameters (2*N*_*e*_s and 2*N*_*e*_*u*) are expected to exhibit similar trajectories for relative changes in allele frequency, genetic load, and fitness components. It thus provides a basis for considering how our models may extrapolate to bottlenecks with much larger *N* than those we have considered.

The validity of using scaled parameters in relation to bottlenecks is examined in Appendix S9, with numerical examples that support its use, at least as a useful approximation (Figures S10 and S11). For example, the results for *s* = 0.5 and *h* = 0.2 with *N* = 50 shown in Figure 2 provide an approximation to the expected behavior in a bottlenecked population of 50 times the size (*N* = 2500) with *s* = 0.01. How well this approach extends to multi-locus systems with linkage remains to be established. In particular, when multiple loci along a chromosome are simulated, the nonlinear relationship between map distance and recombination rate means that recombination cannot in general be rescaled proportionally with *N, s* and *u*. Simulations of multi-locus selection across large genomic regions may therefore be particularly sensitive to the choice of scaling factor (reviewed by Johri et al. 2026), warranting further investigations in relating to the effect of bottlenecks.

If rescaling is valid, at least as an approximation, there is scope for a reduction in the burden of deleterious mutations in bottlenecked populations of sizes in the hundreds or low thousands, provided that they meet the criteria that *h* < ⅓ and 2*Nhs* > 1. These parameters are, of course, difficult to estimate for real data, and variation among studies in bottleneck sizes, times since the bottleneck, and criteria for classifying mutations as deleterious probably explains differences in the conclusions among studies (see Simons and Sella 2016; Robinson et al. 2023). Perhaps counter-intuitively, purging of partially recessive deleterious mutations is most likely to be detected at the genomic level with bottleneck sizes in the thousands rather than the hundreds, as in the example just given. But even with the above example, the expected reduction in burden is likely to be modest (about 8% of the ancestral value) after 750 generations, and a final value of 12%).

### Conclusions

The fate of deleterious mutations in a bottlenecked population of effective size *N*_*e*_ depends directly on the strength of selection and population size. The occurrence of genetic purging at the level of mean allele frequencies, and a temporary recovery in population mean fitness after an initial reduction, require the scaled selection coefficient *γ* = 2*N*_*e*_*s* to be substantially greater than 1, and the dominance coefficient *h* to be less than ⅓. Under these conditions, the inbreeding load *B* relative to the ancestral value may either decrease more rapidly or more slowly than is predicted by 1 – F, where *F* is the neutral inbreeding coefficient of the population. If these conditions on *γ* and *h* are not met, mean allele frequencies stay approximately constant in the short term, mean fitness decreases over time, and *B* changes in proportion to 1 – *F*. Even more stringent conditions are required for the population mean fitness to increase above the ancestral value. The conditions described here represent expectations across many independent stochastic realizations, so that the behavior of an individual population may differ substantially from these expectations, although Figure 5 suggests that the bulk of replicate populations have fairly similar trajectories of mean fitness and inbreeding depression in the multi-locus case.

For very small populations of the kind used in laboratory studies, the fact that the DFE of deleterious mutations is dominated by weakly selected variants implies that most mutations behave nearly neutrally after a bottleneck, so that a recovery of mean fitness recovery is unlikely to be detectable. However, the fact that strongly deleterious and highly recessive mutations contribute disproportionately to *B* means that they could result in a larger reduction in *B* over time than is predicted by 1 – *F*. In bottlenecked natural populations of several hundreds or thousands of individuals, selection should remain effective against a broader range of deleterious mutations, making reductions in their frequencies potentially detectable.

## Data availability

The Fortran 90 source codes and outputs used for the single-locus simulations can be found in https://gitlab.com/elcortegano/oneLocusSim. The SLiM (v4) code used for multi-locus simulations can be found in https://gitlab.com/elcortegano/SLiM_recipes.

## Acknowledgments

Analyses performed here made use of the high-performance computing resources at the Ashworth Compute Co-operative Cluster (AC3) at the Institute of Ecology and Evolution of the University of Edinburgh. We thank two reviewers and Denis Roze for their comments on the previous version of this paper, which helped greatly to improve it.

## Appendix

### 1. The linear operator (LO) method: the case of strong selection

As described in the main text, recursion relations for the population statistics in a bottlenecked population can be found using Ohta’s and Kimura’s linear differential operator method for determining the expectation of a function of a stochastic variable that obeys the diffusion equation approximation (Ohta and Kimura 1969).

Equations (9) imply that, for a function *f*(*g*)of allele frequency *q* we have:

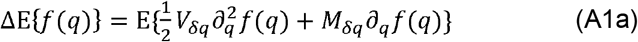

In the case of a single locus in a Wright-Fisher population of size *N*, with the mutation/selection/drift model used in the main text we have:

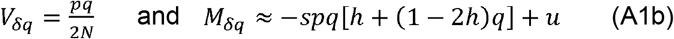

Let the *i*th moment about zero of the distribution of *q* be *M*_*i*_, where M_0_ = 1 and 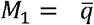; we have 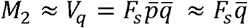, assuming 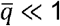 (the rare-allele assumption). Consider *M*_1_, for which *f*(*q*) = *q*. The first and second derivatives of *f*(*q*) are 1 and zero, respectively, so that we only need to find the expectation of 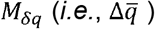. Using Equations (A1), we obtain the following expression:

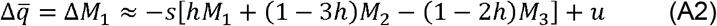

In addition:

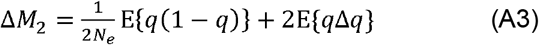

After substituting the expression for 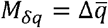 into Equation (A3), this expression yields:

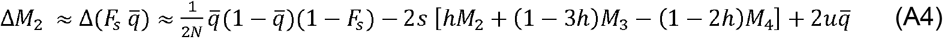

Further progress is hampered by the well-known fact that the non-zero terms involving *q*^2^ in M_δq_ when selection acts mean that a closed system of recursion relations for the moments cannot be obtained (Evans et al. 2007; Gravel 2016; Ragsdale 2022), although such a set can be obtained in the neutral case (Crow and Kimura 1970, pp.331-337). Here we use approximate expressions for the higher moments, which are different for the strong and weak selection cases. In the rest of this section, strong selection (2N*s* ≧ 1for the bottlenecked population) is assumed.

#### Case of *h* > 0 (Method 2a)

We use the following heuristic approach when *h* > 0. As noted in the section *Population genetic statistics for the ancestral population*, the stationary distribution of *q* is approximated by a gamma distribution if *N*_*e*0_*s* is ≫ 1. If it can be assumed that a perturbed distribution rapidly approaches a gamma distribution with shape and scale parameters *α* and *β*, by using the standard formulae for the moments of the gamma distribution we can write 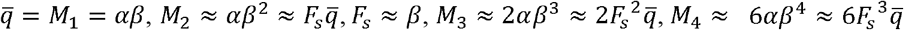, assuming that higher-order powers of *M*1can be neglected. We then obtain the equivalents of Equations (A2) and (A4):

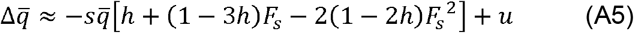

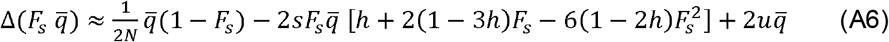

In addition:

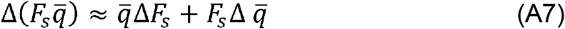

Subtracting 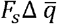 from Equation (A6) and dividing the result by 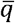 yields:

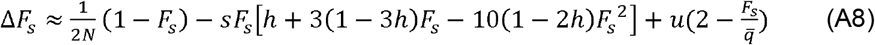

Equations (A5) and (A8) provide a complete, but approximate, description of the dynamics of 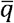 and *F*_*s*_. Their non-linearity means that numerical iteration is the most practical means of generating results, except for the equilibrium state, when both 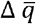 and Δ*F*_*s*_ are equal to zero. Equation (A6) yields the following cubic equation for the equilibrium value of *F*_*s*_, ignoring the term in *u*, which is likely to be very small:

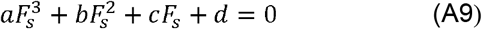

where:

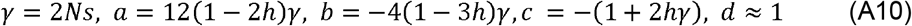

This equation can be solved either by using the standard algebraic solution for a cubic equation, or numerically by Newton-Raphson iteration. In the case of strong selection, *F*_*S*_ is small when *N* is small, so the term involving 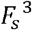 can be neglected, and we obtain the equilibrium expression:

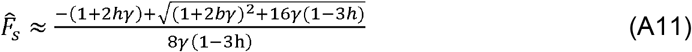

Equation (A5) yields the following result for equilibrium mean *q*:

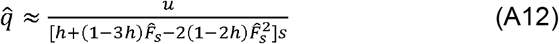

#### Case of complete recessivity (Method 2b)

A similar, but more complex, argument applies when *h* = 0. In this case, the equilibrium distribution of *q* takes the following approximate form, ignoring the small contribution from back mutations (Nei 1968):

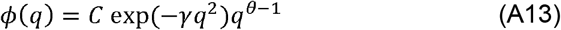

where *θ* = 4*N*_*e*_*u*.

With sufficiently strong selection, the integral of *ϕ* (*q*) between 0 and 1 can be approximated by the integral between 0 and infinity. This integral can be simplified by the substitution 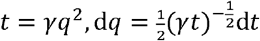, allowing *C* to be obtained:

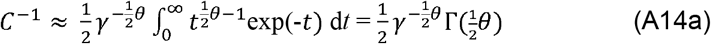

Since θ is small, 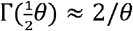, so that:

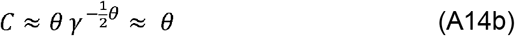

The moments of *ϕ* (*q*) can be obtained by neglecting the small quantity *θ* in the expression for *ϕ*(*q*), using integrals of the following form:

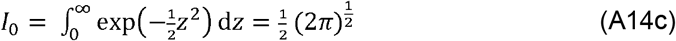

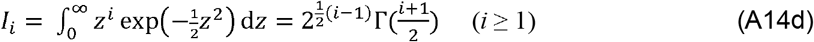

Equation (A14d) yields the following well-known formulae:

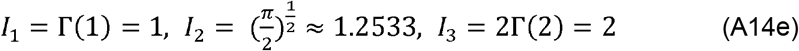

Using the substitution 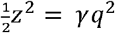, we obtain the following approximate expression for the *i*th moment:

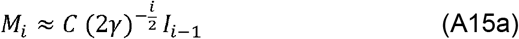

Equation (A14b) implies that:

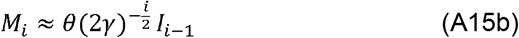

so that:

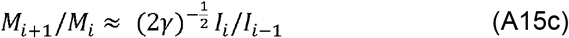

It is easily verified that Equations (A14e) and (A15) can be used to obtain the approximate expressions for the equilibrium values of *M*_1_ and *M*_2_ given by Nei (1968), as well expressions for the higher moments. To deal with the non-equilibrium case, the assumption about the relations between successive moments that was used for the case of *h* > 0 can be applied, which is equivalent to using Equation (A15c) to obtaining *M*_3_ and *M*_4_ in terms of *M*_1_ and *M*_2_, *i*.*e*., 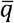 and 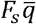. This procedure yields the expressions:

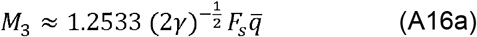

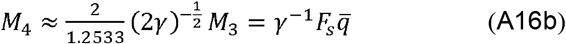

These expressions can be substituted into the basic equations for 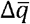 and 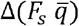 in the same way as in the *h* > 0 case, yielding expressions for 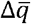 and δ*F*_*s*_. In this case, the terms in *h* vanish, so that the resulting expressions are simpler in form than when *h* > 0. Writing 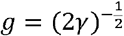, the approximate recursion equations are as follows:

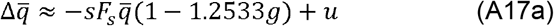

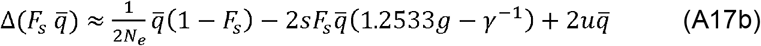

The corresponding equilibrium equations are:

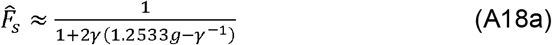

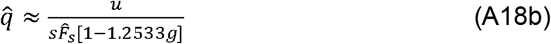

#### Initial conditions

In this way, a set of closed recursion equations for *M*_1_ and *M*_2_ can be obtained and used to determine the population statistics of interest for each generation by numerical iteration, given the initial values of *M*_1_ and *M*_2_. For partially recessive mutations With *N*_*e*0*s*_ ≪ 1, the expected value of *q* given *q*^∗^ = *u* /(*hs*) can be used. For recessive mutations, the initial expected value 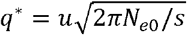 is used (Nei 1968). If it assumed that *Nq*^∗^≪ 1, so that an A_2_ allele is either absent or is present as a single copy, the frequency of A_2_ conditional on survival of the founding event is *q*_0_ = 1/(2*N*), so that 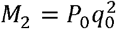, where 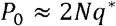 is the probability of survival of A_2_. Similarly, the initial values of *M*_*i*_ for *i* > 2 are given by 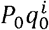.

### 2. The linear operator method with weak selection (Methods 2c and 2d)

In this section, the case of weak selection (*Ns* ≤ 1) is analysed, for which drift can cause *q* to reach non-negligible values, even over the limited timescale of laboratory studies of bottlenecked populations. This means that purging in terms of reduced frequencies of deleterious mutations is unlikely to occur, and the mean fitness of the population is likely to be reduced if deleterious mutations are recessive or partially recessive. The linear operator method can be used in this case, by assuming that changes in the higher order moments of the distribution follow the known recursion relations for neutral mutations, but allowing the recursions for the first and second moments to include the effect of selection and mutation.

With weak selection, Equation (A2) can still be used to determine the recursion for *M*_1_. We also have:

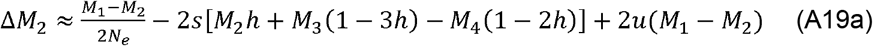

Using Equations (7.4.13 and 7.4.14) of Crow and Kimura (1970, p.234), the neutral recursions for *M*_3_ and *M*_4_ are:

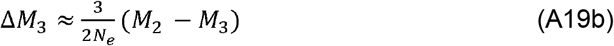

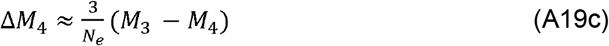

Numerical iteration of this system of equations allows all the properties of interest for a given set of parameters to be determined for each generation of the bottlenecked population (Method 2c).

The process can be further approximated by assuming that 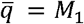 remains constant (as is expected under complete neutrality) and the fixation index obeys the neutral recursion (Method 2d):

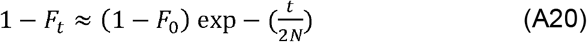

For non-recessive mutations, the genetic load at time *t* relative to its value for the ancestral population then approximated by:

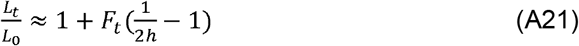

The relative inbreeding load is approximated by:

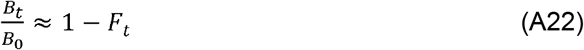

Ultimately, the stationary distribution of *q* will be approached; modeling of this situation must take reverse mutations into account (Kimura *et al*. 1963; Charlesworth 2018). If *γ* is close to zero, the stationary distribution of *q* will be close to that under neutrality. If reverse mutations from A_2_ to A_1_ occur at rate *v*, this is a beta distribution with mean 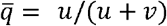 and fixation index given by 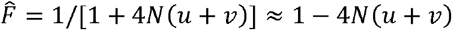 (Wright 1931).

