## Supplementary Material for "The effect of a reduction in population size on mean fitness and inbreeding depression"

**Appendix S1.** Approximations for the trajectories of $q$ and $F_s$

- **Section S1.1.** Approximate allele frequency dynamics for the case of $h > 0$
  - **Equations S1-S3**
- **Section S1.2.** Approximate dynamics of $F_s$ with strong selection and $h > 0$
  - **Equations S4-S7**
- **Section S1.3.** Allele frequency dynamics for the case of $h = 0$
  - **Equations S8-S9**
- **Section S1.4.** The contribution of mutations to allele frequency change with $h = 0$
  - **Equations S10**

**Appendix S2.** Approximation for the expected time ($t_c$) at which purging of deleterious mutations becomes manifest at the level of population mean fitness

- **Section S2.1.** Analytical results
  - **Equations S11-S13**
- **Section S2.2.** Numerical results for the approximation for $t_c$ with strong selection
  - **Figure S1.** Number of generations for purging to become manifest

**Appendix S3.** Simulation parameters

- **Table S1.** Selection models

**Appendix S4.** Extended results for single-locus simulations under strong selection

- **Figure S2.** Results for single-locus simulations and approximations with $h = 0.1$

**Appendix S5.** Extended results for single-locus simulations under weak selection

- **Figure S3.** Results for single-locus simulations and approximations with $h = 0.1$
- **Figure S4.** Results for single-locus simulations and approximations with $h = 0.2$
- **Figure S5.** Results for single-locus simulations and approximations with $h = 0$

**Appendix S6.** Multi-locus simulations

- **Figure S6.** Change in $\bar{L}_r$ over the generations
- **Figure S7.** Change in $\bar{B}_r$ over the generations
- **Figure S8.** Deviations between observed and expected critical purging times

**Appendix S7.** The conditions for the equilibrium deleterious allele frequency and genetic load to be less than their ancestral values

- **Section S7.1.** Conditions on equilibrium $\bar{q}_r$
  - **Equations S14**
- **Section S7.2.** Conditions on equilibrium $\bar{L}_r$
  - **Equations S15-S17**

**Appendix S8.** The limit of $t_c$ as a function of $N_e$

- **Figure S9.** Influence of $s$ and $h$ on $t_c$

**Appendix S9.** Scaling of $s$ and $u$ by $2N_e$

- **Equation S18**
- **Figure S10.** Results of parameter rescaling under relatively strong selection
- **Figure S11.** Results of parameter rescaling under relatively weak selection

**Appendix S10.** Relation between $\bar{q}_r$ and the ratio of diversity at selected sites to diversity at neutral sites

- **Equations S19-S21**

**References**

#### Appendix S1. Approximations for the trajectories of *q* and $F_s$

##### 1.1. Approximate allele frequency dynamics for the case of *h* > 0

We use the convention that $t = 1$ is the generation immediately following the bottleneck, for which $F_{s\,1} = 1/(2N)$. $F_{s\,0}$ for the ancestral population can be set to zero as a good approximation, given the assumption that $N_0 hs \gg 1$. Using Equation (5b) of the main text, together with Equation (S6c) for $F_{s\,t}$ derived in section 1.2 below, we obtain the following approximation for the dynamics of the mean allele frequency:

$$\frac{\mathrm{d}\ln(\bar{q}_t)}{\mathrm{d}t} \approx -s\{(1-3h)[\hat{F}_s + (F_{s\,0} - \hat{F}_s)\exp\left(-\frac{(1+2h\gamma)t}{2N}\right)]\} \qquad \text{(S1)}$$

where the equilibrium value of $F_{s\,t}$ is given by $\hat{F}_s \approx 1/(1+2h\gamma)$. Integration of this expression with respect to *t*, and writing $\bar{q}_{r\,t} = \bar{q}_t/q^*$ for the ratio of the value of mean of *q* to its equilibrium value $q^*$, we obtain:

$$\ln(\bar{q}_{rt}) \approx -s\{(1-3h)\hat{F}_s t + 2N(1-3h)\left(F_{s\,0} - \hat{F}_s\right)\hat{F}_s[1 - \exp\left(-\frac{(1+2h\gamma)t}{2N}\right)]\} \qquad \text{(S2)}$$

so that

$$\bar{q}_{r\,t} \approx \exp -s\{(1-3h)\hat{F}_s t + 2N(1-3h)\left(F_{s\,0} - \hat{F}_s\right)\hat{F}_s\left[1 - \exp\left(-\frac{(1+2h\gamma)t}{2N}\right)\right]\} \qquad \text{(S3)}$$

This expression is included in the main text as Equation (6).

**1.2. Approximate dynamics of $F_s$ with strong selection and $h$ > 0**

An approximation for the recursion for $F_s$ with selection against non-recessive mutations can be obtained as follows. If the change in allele frequency per generation is small and the value of $\bar{q}$ in generation $t$, $\bar{q}_t$, is also small, then the change in the value of $F_s$ between generation $t$ and $t$ – 1, $\Delta F_t$, is approximately $\Delta M_{2\,t}/\ \bar{q}_t$, where $\Delta M_{2\,t}$ is the change in the second moment of $q$ between generations $t$ and $t$ – 1. Neglecting higher moments in the expression for $\Delta M_{2\,t}$ in Equation (A4), the following approximation for $\Delta F_t$ is obtained, assuming that $2Nhs > 1$:

$$\Delta F_{s\,t} \approx \frac{1}{2N} - (\frac{1+2h\gamma}{2N})F_{s\,t-1} \qquad \text{(S4)}$$

where $\gamma = 2Ns$. This expression yields the approximate equilibrium value of $F_s$ as:

$$\hat{F}_s \approx \frac{1}{1+2h\gamma} \qquad \text{(S5)}$$

This corresponds to the well-known result obtained when approximating the stationary distribution of $q$ under mutation and selection by assuming only small deviations from the mean of $q$ (e.g., Charlesworth and Charlesworth 2010, pp.354-355).

Equation (S4) can then be rewritten as follows:

$$\Delta(F_{s\,t} - \hat{F}_s) \approx \frac{1}{2N} - \left(\frac{1+2h\gamma}{2N}\right)F_{s\,t-1} = -\left(\frac{1+2h\gamma}{2N}\right)(F_{s\,t-1} - \hat{F}_s) \qquad \text{(S6a)}$$

$$(F_{s\ t} - \hat{F}_s) \approx (F_{s\,0} - \hat{F}_s)\left[1 - \left(\frac{1+2h\gamma}{2N}\right)\right]^t \qquad \text{(S6b)}$$

where $F_{s\,0}$ is the fixation index for the ancestral population; $F_{s\,0} \approx 0$ under the assumptions made here.

We thus have:

$$F_{s\ t} \approx \hat{F}_s + (F_{s\,0} - \hat{F}_s)\left[1 - \left(\frac{1+2h\gamma}{2N}\right)\right]^t \approx \hat{F}_s + (F_{s\,0} - \hat{F}_s)\exp\left(-t\frac{1+2h\gamma}{2N}\right) \qquad \text{(S6c)}$$

If $t$ is small compared with $2N$, the exponential term can be approximated by $1 - T(1 + 2h\gamma)$, where $T = 1/2N$. This implies that the change in $F_s$ per generation is given by:

$$\Delta F_{s\,t} \approx (\hat{F}_s - F_{s\,0})\frac{(1+2h\gamma)}{2N} \approx \frac{1}{2N} \qquad \text{(S7)}$$

where the final approximation is valid if $F_{s0} \ll \hat{F}_s$. Thus, for the early generations after the bottleneck, $\Delta F_{s\,t}$ is likely to be close to the neutral value, independently of the strength of selection. This result should also hold for the case of completely recessive mutations ($h$ = 0), since selection is ineffective against rare recessive mutations.

**1.3. Allele frequency dynamics for the case of *h* = 0**

The approach of section 1.1 above cannot be used if $h = 0$. Instead, we approximate Equation (7) of the main text by assuming that $N$ is sufficiently small that $F_{s\,1} \gg \bar{q}$. In addition, the neutral recursion for *F* must be used, as there is no useful approximation for the effect of selection when $h = 0$. Ignoring the mutational contribution, which is shown in section 1.4 below to be negligible when $h = 0$, we have:

$$\frac{\mathrm{d}\ln(\bar{q})}{\mathrm{d}t} \approx -sF \approx -s\left[1 - \exp\left(-\frac{t}{2N}\right)\right] \qquad \text{(S8a)}$$

$$\ln(\bar{q}_{r\,t}) \approx -s\left\{t - (2N)\left[1 - \exp\left(-\frac{t}{2N}\right)\right]\right\} \qquad \text{(S8b)}$$

so that:

$$\bar{q}_{r\,t} \approx \exp -s\left\{t - (2N)\left[1 - \exp\left(-\frac{t}{2N}\right)\right]\right\} \qquad \text{(S8c)}$$

For sufficiently small *F*, this can be further approximated using the first three terms in the series expansion of $\exp(x)$, yielding:

$$\bar{q}_{r\,t} \approx \exp\left(-\frac{st^2}{4N}\right) \qquad \text{(S8d)}$$

The relative value of the inbreeding load is given by:

$$\bar{B}_{r\,t} \approx \bar{q}_{r\,t}(1 - F_{s\,t}) \qquad \text{(S9a)}$$

Using Equation (S8d) and approximating $1 – F_{s\,t}$ by the neutral expression $\exp\left(-\frac{t}{2N}\right)$, this expression can be further approximated by:

$$\bar{B}_{r\,t} \approx \exp\left(-\frac{2t + st^2}{4N}\right) \qquad \text{(S9b)}$$

Equations (S8c) and the expression for neutral *F* can be used to determine the relative genetic load at time *t* by substitution into Equation (1) of the main text.

**1.4. The contribution of mutations to allele frequency change with *h* = 0**

The justification for ignoring the mutational term in the fully recessive case is as follows. If $F \gg \bar{q}$, the initial population is at the deterministic equilibrium under mutation and selection with $q^* \approx \sqrt{u/s}$ and $\bar{q}$ is close to $q^*$, we can write:

$$\frac{\Delta\bar{q}}{\bar{q}} \approx -sF + \frac{u}{\bar{q}} \approx -s(T - \frac{u}{q^*}) = -s(T - \sqrt{us}) = -s(T - \sqrt{u/s}) \qquad \text{(S10a)}$$

where $T = t/(2N)$. For realistic values of $T$, $u$ and $s$, the term in $\sqrt{u/s}$ can be ignored.

However, in reality there is a wide probability distribution of $q$ for completely recessive mutations, even in very large populations, with mean $q^* \approx u\sqrt{2\pi N_{e0}/s} \ll \sqrt{u/s}$ and variance $V_{q0} \approx u/s$ (Nei 1968), where $N_{e0}$ is the effective size of the ancestral population. The genetic load for the initial population is thus approximately equal to $u$, provided that $q^* \ll 1$, which is the same as the infinite population value.

In this case, Equation (S10a) is replaced by:

$$\frac{\Delta\bar{q}}{\bar{q}} \approx -sF + \frac{u}{q^*} \approx -sT + \frac{1}{\sqrt{2\pi N_{e0}/s}} \qquad \text{(S10b)}$$

Since $2\pi N_{e0} >> 1$ and $s \leq 1$, the second term in this equation can again be ignored.

**Appendix S2. Approximation for the expected time ($t_c$) at which purging of deleterious mutations becomes manifest at the level of the population mean fitness**

#### 2.1. Analytical results

Here we obtain an approximation for the minimum expected number of generations required for genetic purging to become manifest at the level of the population mean fitness, *i.e.*, the "critical time for purging", $t_c$ (López-Cortegano 2020). Strongly selected mutations ($Ns \gg 1$) are assumed for the purpose of this calculation. Formally, $t_c$ is the time at which the derivative of expected mean fitness (or genetic load) with respect to time is zero, provided that the initial trend of mean fitness is downwards and allele frequency change is sufficiently slow that time can be treated as a continuous variable.

In order to obtain tractable expressions, we approximate $F_s$ by equating it to the neutral fixation index, $F$. The fact that $\bar{w}$ can be written as a function of $\bar{q}$ and $F$ implies that it has the following derivative with respect to time:

$$\frac{\partial \bar{w}}{\partial t} \approx -s\frac{\mathrm{d}\bar{q}}{\mathrm{d}t}[2h + (1-2h)F] - s\bar{q}(1-2h)\frac{\mathrm{d}F}{\mathrm{d}t} \qquad \text{(S11a)}$$

Writing $\frac{\mathrm{d}\bar{q}}{\mathrm{d}t} \approx \Delta\bar{q}$ and $\frac{\mathrm{d}F}{\mathrm{d}t} \approx \Delta F$, we have:

$$\frac{\partial \bar{w}}{\partial t} \approx -s\Delta\bar{q}[2h + (1-2h)F] - s\bar{q}(1-2h)\Delta F \qquad \text{(S11b)}$$

We also have $\Delta F = (1-F)/2N \approx 1/2N$, provided that $t$ is sufficiently small relative to $2N$. This implies that $F \approx T$, where $T = t/(2N)$ is the time in units of $2N$ generations, with $t = 1$ corresponding to the first generation of the bottlenecked population. For small $T$, this is a good approximation even in the presence of selection, as is shown by Equation (S7). For the case of $h > 0$, using Equation (4) of the main text we can write:

$$\frac{\Delta\bar{q}}{\bar{q}} \approx -s\left[h + (1-3h)T - \frac{u}{s\bar{q}}\right] \qquad \text{(S11c)}$$

For small $t$ and $h > 0$, $\bar{q} \approx q^*$, and $q^* = u/(hs)$ (see the subsection of the main text *Population genetic statistics for the ancestral population*). The terms in $h$ and $u/s\bar{q}$ in this expression thus approximately cancel each other, and Equation (S11b) yields:

$$\frac{1}{\bar{q}}\ \frac{\partial \bar{w}}{\partial t} \approx\ (1-3h)[2h+(1-2h)T]Ts^2 - \frac{(1-2h)s}{2N} \qquad \text{(S11d)}$$

The positive term in $s^2$ corresponds to the effect on expected mean fitness of the decrease in expected allele frequency under selection, minus the effect of mutation, and the term – (1 – 2$h$)$s$/(2$N$) corresponds to the effect of the increase in $F$ over one generation. A slightly different approach to justifying the neglect of the effect of mutation is needed for the case of complete recessivity ($h$ = 0) (see section 1.4 above).

By definition, $t_c$ corresponds to the value of $t$ for which this expression is zero, yielding a quadratic equation for the critical value of $T_c = t_c/(2N)$. Writing $\gamma = 2Ns$, this equation can be written as:

$$(1-2h)(1-3h)T_c^2 + 2h(1-3h)T_c - (1-2h)\gamma^{-1}\ \approx 0 \qquad \text{(S12)}$$

If $h$ < ½, as is required for inbreeding depression, this equation has the following solution:

$$T_c \approx \frac{-h+\sqrt{h^2+(1-2h)^2[(1-3h)\gamma]^{-1}}}{(1-2h)} \qquad \text{(S13a)}$$

The corresponding time in generations is $t_c = 2NT_c$. This equation has a meaningful solution with $T_c > 0$ only if $h < \frac{1}{3}$; otherwise, the expected mean fitness will always decline with time or remain approximately stationary. This equation is included in the main text as Equation (8).

When $h$ is small in relation to $\gamma$ and mutation is ignored, we have:

$$T_c \approx \sqrt{1/\gamma(1-3h)} \qquad \text{(S13b)}$$

or equivalently:

$$t_c \approx \sqrt{\frac{2N}{s(1-3h)}} \qquad \text{(S13c)}$$

If $h > 0$, this expression is recovered if the linear term in $T_c$ in Equation (S12) is neglected. In practice, the second approximation does not yield satisfactory results when compared with single-locus simulations unless $h \approx 0$; otherwise, it tends to overestimate $T_c$ (see the numerical results presented in the main text).

##### 2.2. Numerical results for the approximation for $t_c$ with strong selection

Figure S1 shows some representative examples of $t_c$ as a function of $N$ and $s$, using Equation (S13a). Here $h < ⅓$, so that the mean allele frequency $\bar{q}$ is expected to decrease over time, *i.e.*, there is purging. It should be noted that these approximations assume $Ns \gg 1$; some values in the figure are outside this range and are included only for purposes of illustration. The strength of selection is an important determinant of $t_c$, with more deleterious mutations reaching $t_c$ faster than less deleterious ones; this difference is larger for larger $N$. However, although small populations are expected to reach the point of fitness recovery faster than large populations, this comes with the cost of higher rates of genetic drift, as measured by the neutral fixation index $F$ (Figure S1B).

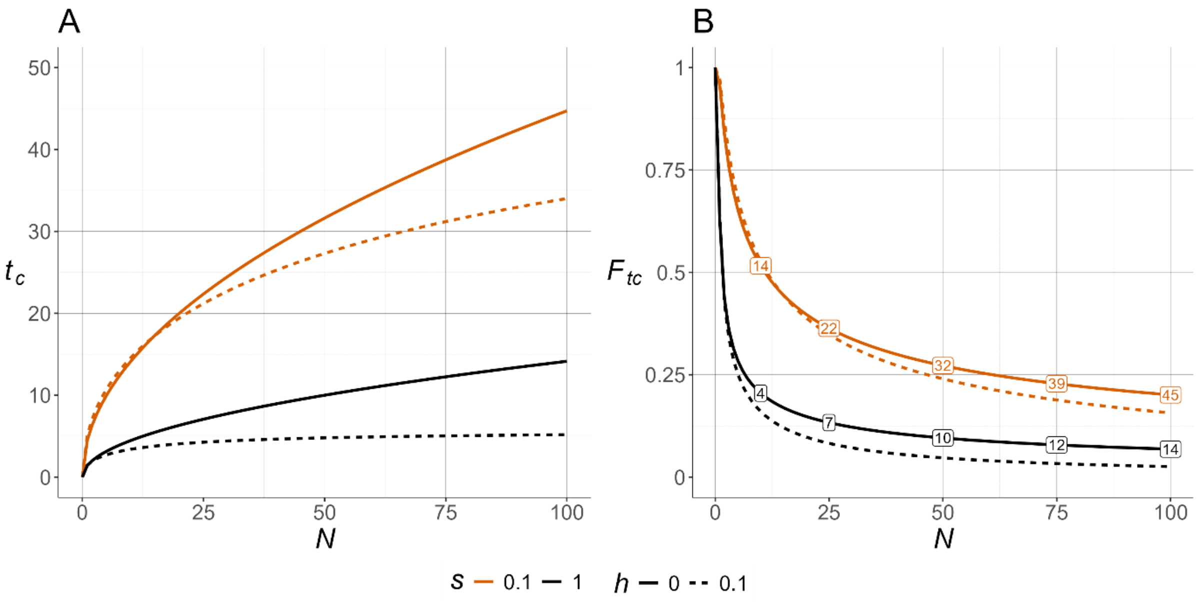


Figure S1. Number of generations for purging to become manifest ($t_c$). A) Approximate theoretical values of $t_c$ as functions of the effective population size ($N$), for different values of the selection coefficient ($s$) and the degree of dominance ($h$), using Equation (S13a). B) The neutral fixation index ($F$) at generation $t_c$ as a function of $N$, for different values of $s$ and $h$. The numbers inset into panel B show some representative approximate values of $t_c$ for the corresponding $F$ values.

#### Appendix S3. Simulation parameters

In all simulations, the base population reached a stable inbreeding load $B \approx 4.5$ after 5,000 generations. To attain this target value, we adjusted the mutation rate ($u$) for each selection model (Table S1).

**Table S1.** Selection models. Mutation rate ($u$) used for selection models with different values of selection coefficient ($s$) and degree of dominance ($h$).

| $s$ | $h$ = 0 | $h$ = 0.1 | $h$ = 0.2 |
|---|---|---|---|
| 0.01 | $2.238 \times 10^{-8}$ | $4.229 \times 10^{-8}$ | $7.759 \times 10^{-8}$ |
| 0.1 | $7.887 \times 10^{-9}$ | $2.902 \times 10^{-8}$ | $7.030 \times 10^{-8}$ |
| 0.2 | $5.694 \times 10^{-9}$ | $2.740 \times 10^{-8}$ | $6.879 \times 10^{-8}$ |
| 0.3 | $4.638 \times 10^{-9}$ | $2.688 \times 10^{-8}$ | $6.863 \times 10^{-8}$ |
| 0.4 | $4.019 \times 10^{-9}$ | $2.659 \times 10^{-8}$ | $6.861 \times 10^{-8}$ |
| 0.5 | $3.600 \times 10^{-9}$ | $2.642 \times 10^{-8}$ | $6.852 \times 10^{-8}$ |
| 0.6 | $3.272 \times 10^{-9}$ | $2.619 \times 10^{-8}$ | $6.848 \times 10^{-8}$ |
| 0.7 | $3.038 \times 10^{-9}$ | $2.609 \times 10^{-8}$ | $6.843 \times 10^{-8}$ |
| 0.8 | $2.843 \times 10^{-9}$ | $2.607 \times 10^{-8}$ | $6.834 \times 10^{-8}$ |
| 0.9 | $2.683 \times 10^{-9}$ | $2.599 \times 10^{-8}$ | $6.832 \times 10^{-8}$ |
| 1.0 | $2.533 \times 10^{-9}$ | $2.591 \times 10^{-8}$ | $6.812 \times 10^{-8}$ |

### Appendix S4. Extended results for single-locus simulations under strong selection

Figure S2 shows results analogous to those in Figures 2 and 3, using $h$ = 0.1.

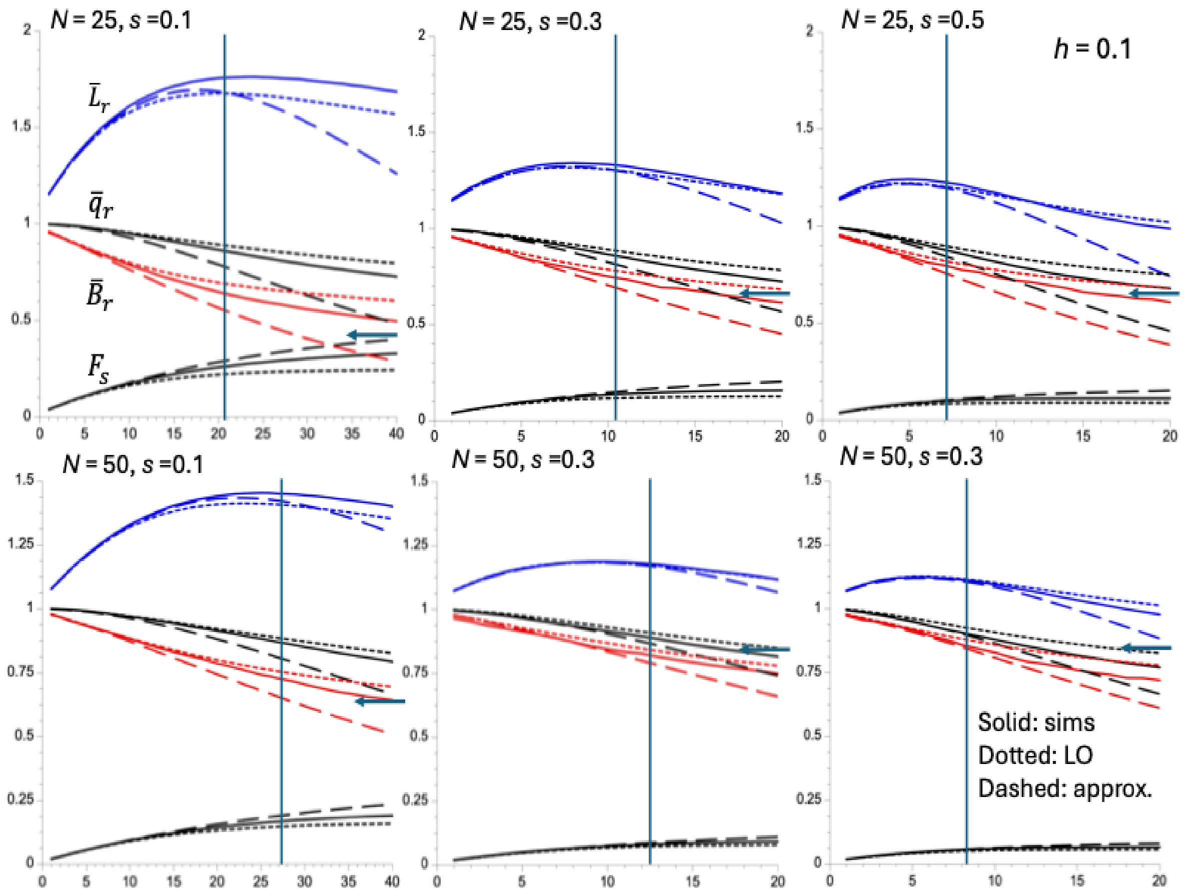


Figure S2. Results of single-locus simulations and approximations with $h$ = 0.1 and strong selection. The solid curves (in order from top to bottom) in each panel show the means of the following statistics over all replicate simulations: the mean genetic load relative to its equilibrium value ($\bar{L}_r$), the mean allele frequency relative to its equilibrium value ($\bar{q}_r$), the mean inbreeding depression relative to its equilibrium value ($\bar{B}_r$) and the mean fixation index at the selected sites ($F_s$). The dotted curves are the predictions based on the linear operator Method 2a, and the dashed curves are based on the approximations given by the trajectories for the mean allele frequency using Method 1a and the corresponding expression for the relative values of the genetic and inbreeding loads. The arrows display the final value of one minus the neutral fixation index, 1 – $F$, which corresponds to $\bar{B}_r$ if there is no effect of selection on allele frequencies. The vertical lines indicate the values of the approximate expected time for purging to be manifested ($t_c$), obtained from Equation (S13a). See the *Methods of Analysis* section for details of the simulation method.

#### Appendix S5. Extended results for single-locus simulations under weak selection

Figures S3-S5 compare results for simulations under weak selection with predictions from the linear operator method.

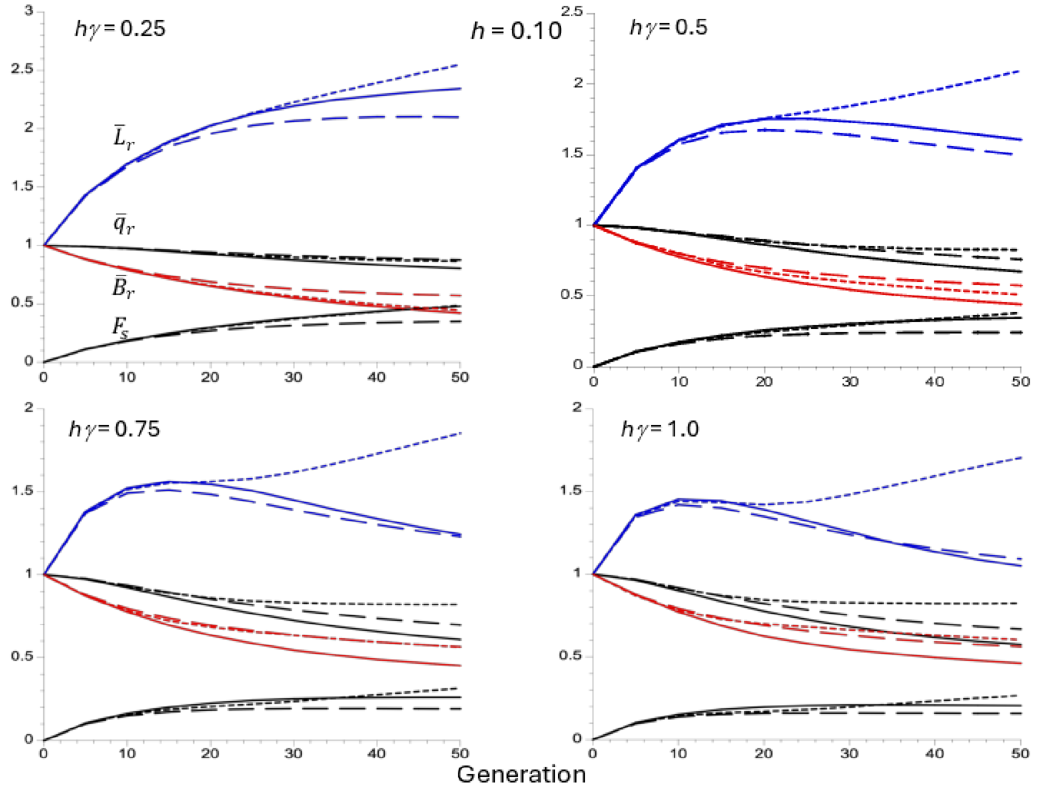


Figure S3. Results for single-locus simulations and linear operator approximations with $h$ = 0.1. The solid curves (in order from top to bottom) in each panel show the means of the following statistics over all replicate simulations: the mean genetic load relative to its equilibrium value ($\bar{L}_r$), the mean allele frequency relative to its equilibrium value ($\bar{q}_r$), the mean inbreeding depression relative to its equilibrium value ($\bar{B}_r$) and the mean fixation index at the selected sites ($F_s$). The dotted curves are the predictions based on the weak selection linear operator Method 2c, and the dashed curves are the predictions based on the strong selection version, Method 2a.

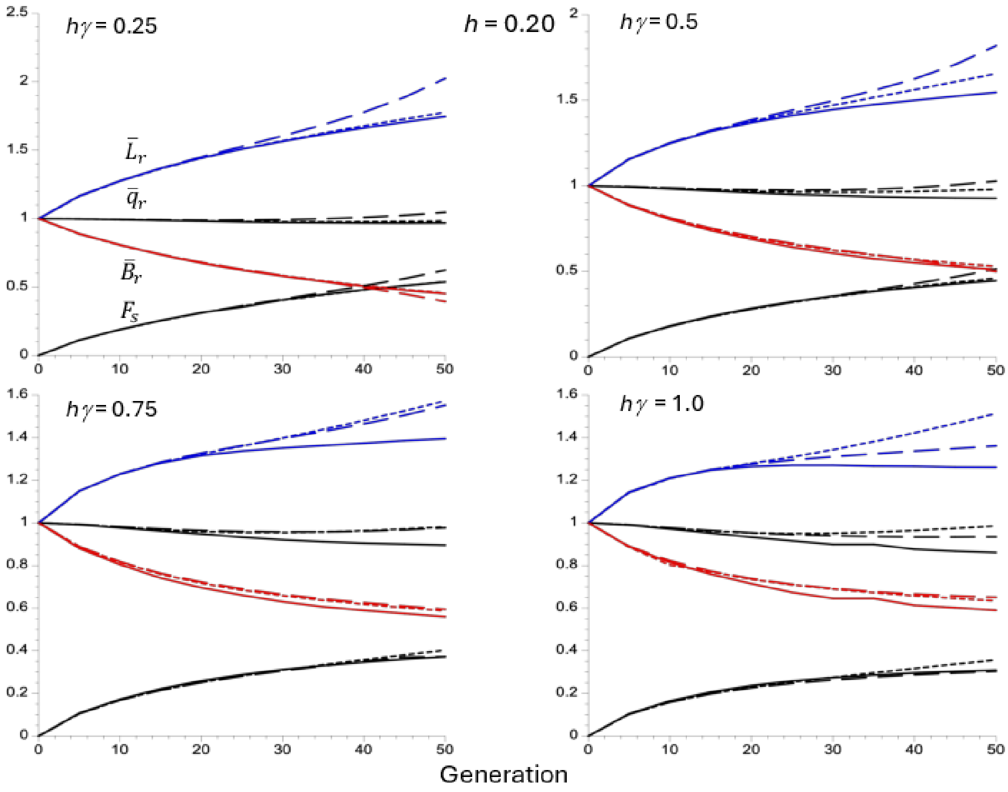


Figure S4. Results for single-locus simulations and linear operator approximations with $h$ = 0.2. The solid curves (in order from top to bottom) in each panel show the means of the following statistics over all replicate simulations: the mean genetic load relative to its equilibrium value ($\bar{L}_r$), the mean allele frequency relative to its equilibrium value ($\bar{q}_r$), the mean inbreeding depression relative to its equilibrium value ($\bar{B}_r$) and the mean fixation index at the selected sites ($F_s$). The dotted curves are the predictions based on the weak selection linear operator Method 2c, and the dashed curves are the predictions based on the strong selection version, Method 2a.

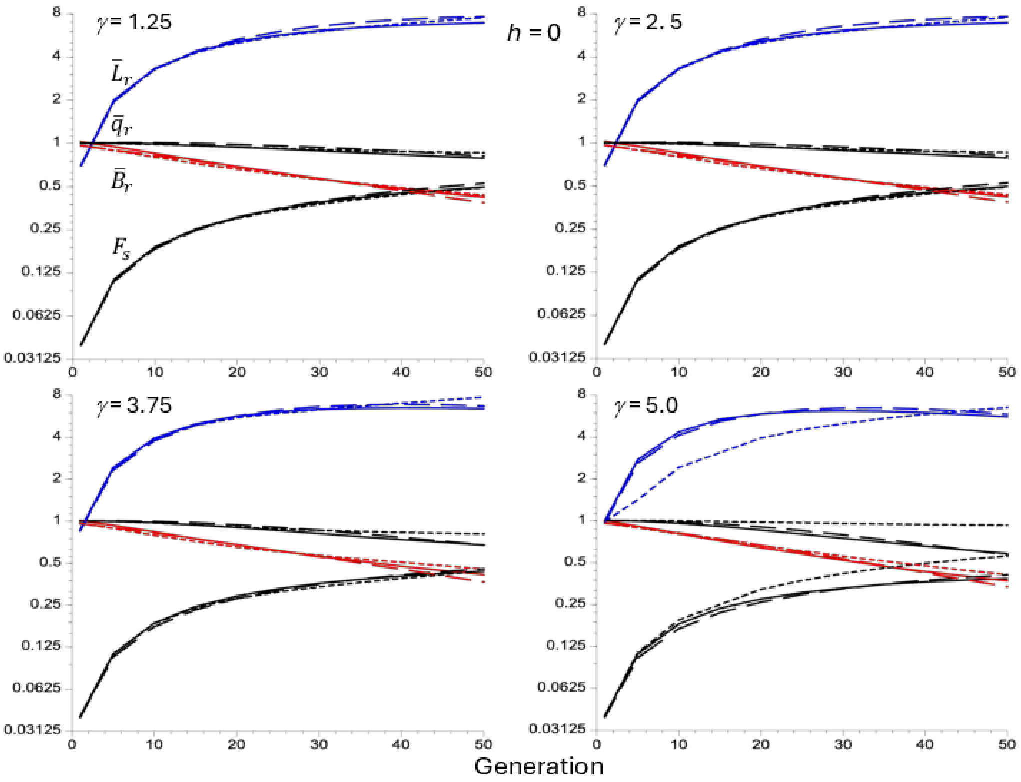


Figure S5. Results for single-locus simulations and linear operator approximations with $h = 0$. The solid curves (in order from top to bottom) in each panel show the means of the following statistics over all replicate simulations: the mean genetic load relative to its equilibrium value ($\bar{L}_r$), the mean allele frequency relative to its equilibrium value ($\bar{q}_r$), the mean inbreeding depression relative to its equilibrium value ($\bar{B}_r$) and the mean fixation index at the selected sites ($F_s$). The dotted curves are the predictions based on the weak selection linear operator Method 2c, and the dashed curves are the predictions based on the strong selection version, Method 2a.

#### Appendix S6. Multi-locus simulations

Multi-locus simulations using run allowed us to examine how associations among loci modify the efficiency and timing of purging (see Methods). Representative examples of generational changes in the relative genetic ($\bar{L}_r$) and inbreeding load ($\bar{B}_r$) are shown in Figures S6 and S7, respectively, comparing the trends of multi-locus simulations with single-locus simulations.

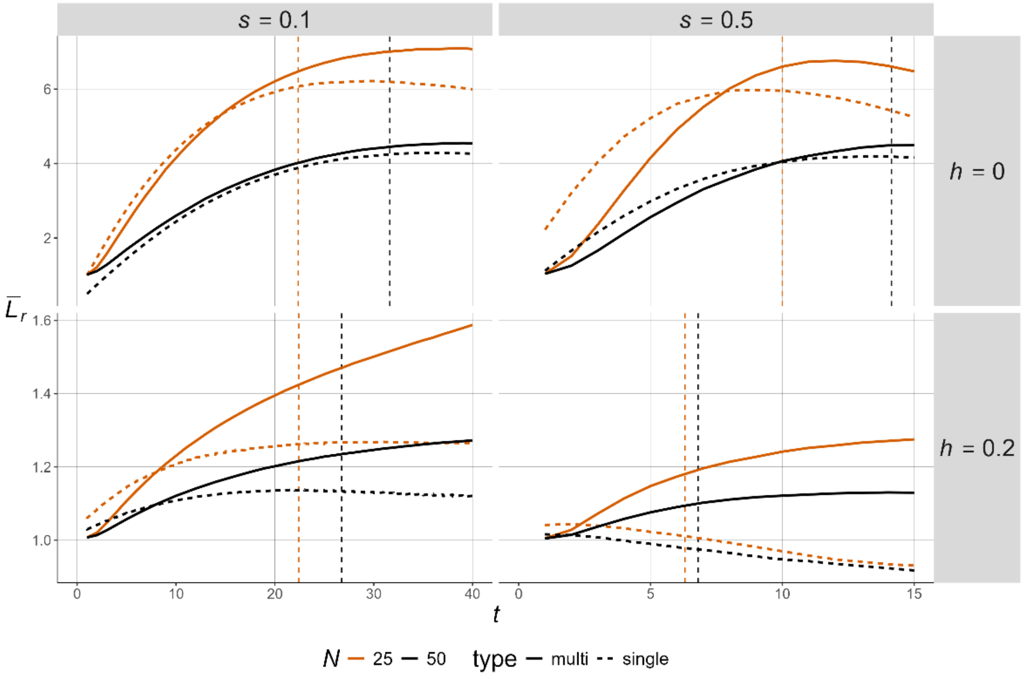


Figure S6. Change in the relative genetic load ($\bar{L}_r$) over generations ($t$). Results of multi-locus with fixed selection coefficients (1,000 simulation replicates) and corresponding single-locus simulations are shown as solid and dashed lines, respectively. Panels from left to right represent increasing selection coefficients, and from top to bottom models with increasing dominance coefficients ($h$). Results for populations of size $N$ = 25 (orange) and $N$ = 50 (black) are shown. The colored, dashed vertical lines mark the analytical value of $t_c$ (Equation (S13a)).

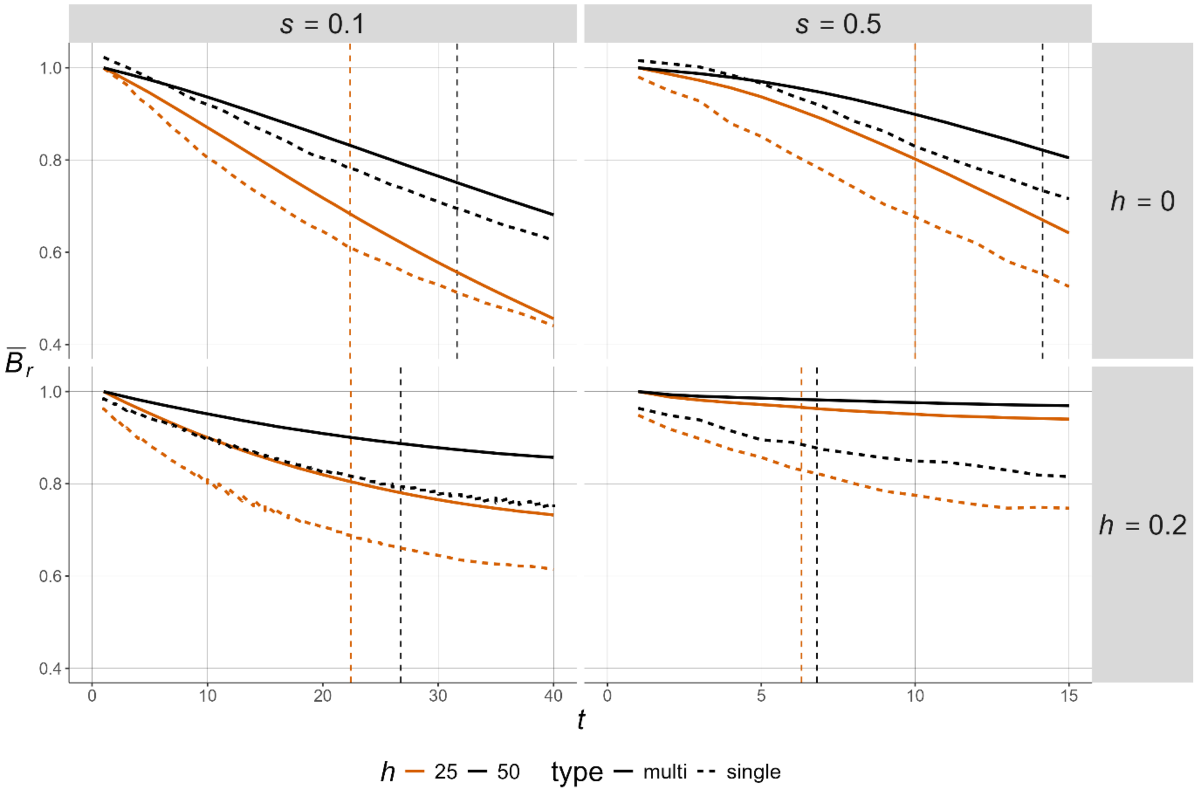


Figure S7. Change in the relative inbreeding load ($\bar{B}_r$) over generations ($t$). Results of multi-locus with fixed selection coefficients (1,000 simulation replicates). Panels from left to right represent increasing selection coefficients, and from top to bottom models with increasing dominance coefficients ($h$). Results for populations of size $N$=25 (orange) and $N$=50 (black) are shown. The colored, dashed vertical lines mark the analytical value of $t_c$ (Equation (S13a)).

To determine the observed critical time of purging, $O(t_c)$, we calculated the mean population mean fitness ($w$) across replicate simulations for each generation and defined $O(t_c)$ as the generation at which $w$ reached its minimum. We then compared $O(t_c)$ with the expectation from Equation (S13c) by calculating $R(t_c) = O(t_c) - E(t_c)$. Figure S8 shows that Equation (S13c) reliably predicts the onset of purging provided that the bottlenecked population has sufficiently large $Ns$, consistent with the common understanding that selection prevails over genetic drift when $N_e s > 1$. The best fit was observed for fully recessive alleles.

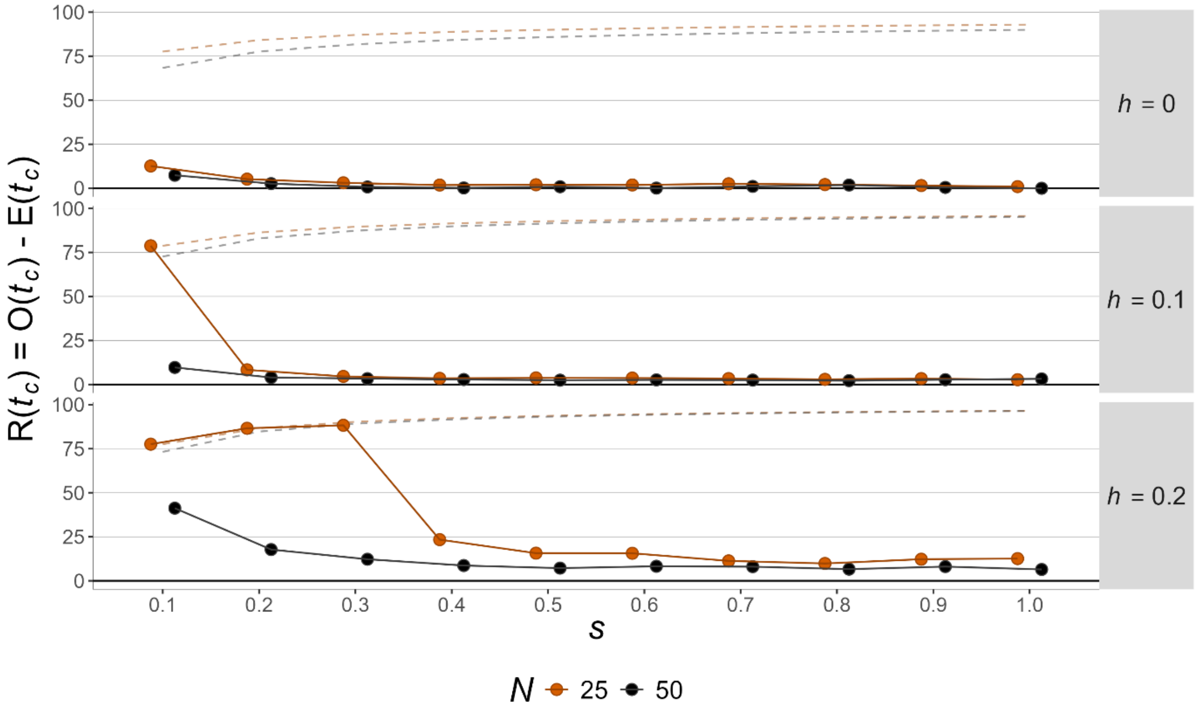


Figure S8. The deviations between observed and expected critical purging times, $R(t_c) = O(t_c) - E(t_c)$, where $O(t_c)$ was calculated using the mean fitness taken over 1,000 replicate simulations for a given population size ($N$), selection coefficient $s$ and dominance coefficient $h$. Only values with $Ns \geq 1$ are shown. Dashed lines represent the values of 100 – E($t_c$), which indicate cases where no purging has occurred. In these cases, the error bars on $R(t_c)$ are necessarily one-sided, since simulations were terminated after 100 generations.

It should be noted that, for simulations with the lowest $Ns$ values, where mutations are nearly neutral, purging effects are negligible, since even weakly deleterious mutations can drift to fixation, resulting in a fitness decline without any recovery, in which case $R(t_c) \approx 100 - E(t_c)$ (Figure S8). However, this process becomes insignificant for most cases with significant selection $Ns \gg 1$. Excluding scenarios with no fitness recovery, the fact that $R(t_c)$ tended to be positive when not overlapping zero suggests that Equation (S13c) (equivalent to Equation (8) of the main text) serves as a lower bound for the time required for purging to be manifest in the multi-locus context (especially for the least recessive case, $h = 0.2$).

### Appendix S7. The conditions for the equilibrium deleterious allele frequency and genetic load to be less than their ancestral values

#### 7.1. Conditions on equilibrium $\bar{q}_r$

Equations (12) of the main text can be combined together to yield the following quadratic expression for the maximal value of *h* that is compatible with a meaningful solution:

$$6\gamma h^2 - (2\gamma + 1)h + 1 < 0 \qquad \text{(S14a)}$$

Solving the corresponding quadratic equation with the right-hand side equal to zero, whose only meaningful solution is the larger root, implies that:

$$h < \frac{(2\gamma+1)+\sqrt{(2\gamma+1)^2-24\gamma}}{12\gamma} \qquad \text{(S14b)}$$

The condition for the discriminant of this expression to be non-negative, so that *h* is a real number, is:

$$4\gamma^2 - 20\gamma + 1 > 0 \qquad \text{(S14c)}$$

This has the solution $\gamma > 4.95$. The corresponding value of *h* is 0.184. Numerical evaluation of Equation (S14b) shows that *h* rapidly approaches its asymptotic value of ⅓ as *γ* increases.

#### 7.2. Conditions on equilibrium $\bar{L}_r$

This analysis also assumes strong selection, and makes use of the results in section 1 of the Appendix for the case of non-recessive mutations (Method 2a). The expected relative genetic load is given by the following expression:

$$\bar{L}_r \approx \frac{[2h+(1-2h)F_S]\bar{q}s}{2q^*hs} = \frac{[2h+(1-2h)F_S]\bar{q}}{2q^*h} \qquad \text{(S15a)}$$

From Equation (A12) and the relation $q^* \approx u/(hs)$, at equilibrium we have:

$$\hat{L}_r \approx \frac{[2h+(1-2h)\hat{F}_S]}{2[h+(1-3h)\hat{F}_S-2(1-2h)\,\hat{F}_S^2]} \qquad \text{(S15b)}$$

We thus have $\hat{L}_r < 1$ if and only if:

$$2h + 2(1-3h)\hat{F}_s - 4(1-2h)\,\hat{F}_s^2 > 2h + (1-2h)\hat{F}_s$$

i.e., $$2(1-3h)\hat{F}_s - 4(1-2h)\,\hat{F}_s^2 > (1-2h)\hat{F}_s$$

This reduces to:

$$4h < 1 - 4(1-2h)\hat{F}_s$$

$$h < \tfrac{1}{4} - (1-2h)\hat{F}_s \qquad \text{(S16)}$$

This expression is included in the main text as Equation (13).

Following the approach used above for equilibrium $\bar{q}_r$, by combining this expression with Equation (12b), we obtain the following quadratic relation:

$$8\gamma h^2 - (2\gamma+4)h + 3 < 0 \qquad \text{(S17a)}$$

This has the solution:

$$h < \frac{(\gamma+2)+\sqrt{(\gamma+2)^2-24\gamma}}{8\gamma} \qquad \text{(S17b)}$$

The condition for the discriminant to be non-negative is:

$$\gamma^2 - 20\gamma + 4 > 0 \qquad \text{(S17c)}$$

This has the solution $\gamma > 19.8$; the corresponding value of $h$ is 0.138.

#### Appendix S8. The limit of $t_c$ as a function of $N_e$

Equations (S13) suggest that $t_c$ takes relatively small values when $h > 0$ and selection is strong ($Ns \gg 1$). Under these conditions, $t_c$ increases with $N$ but quickly approaches an asymptote. Thus, $t_c$ primarily depends on $s$ and $h$. This can be demonstrated by evaluating the limit of Equation (8) as $N \to \infty$, which can be solved as:

$$\lim_{N\to\infty} t_c = \frac{1-2h}{2sh(1-3h)}$$

Figure S9 shows that the limit of $t_c$ as a function of $N$ is just a few tens of generations for a wide range of parameters. Minimum $t_c$ values are observed for $s = 1$ and $h = \frac{3-\sqrt{3}}{6} \approx 0.21$. It can be deduced that for most populations, $t_c$ is relatively small in relation to evolutionary timescales, given strong selection and $h$ sufficiently bounded away from both 0 and ⅓.

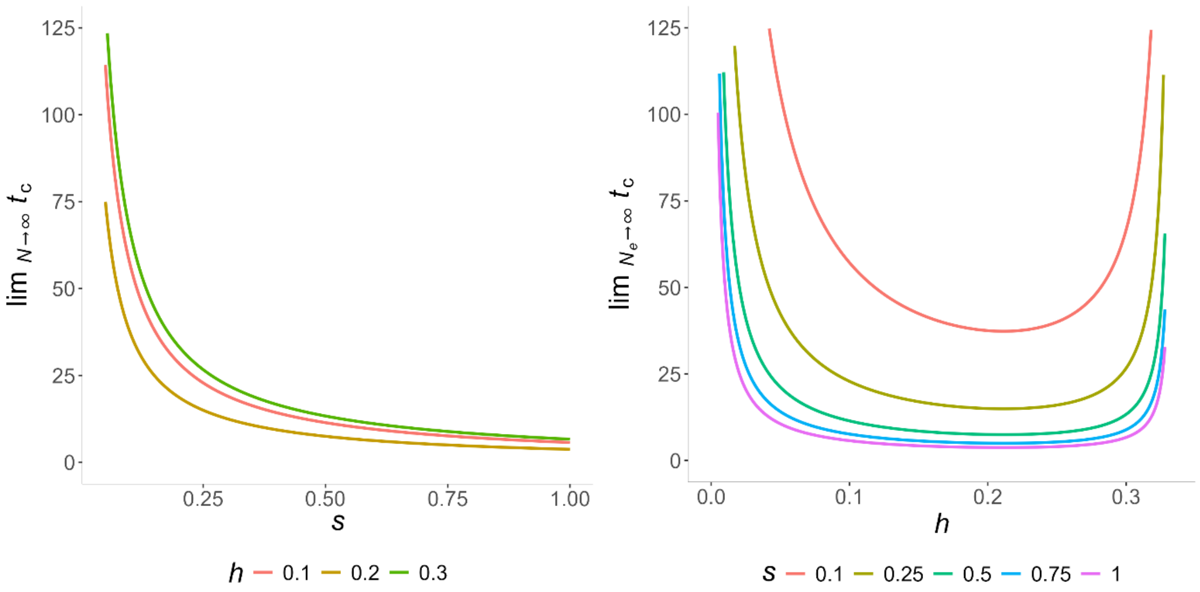


Figure S9. Influence of the selection coefficient ($s$) and dominance coefficient ($h$) on the critical time for purging ($t_c$). A) Change in the limiting value of $t_c$ over values of $s$ (in colours, $h$). B) Change in the limiting value of $t_c$ over values of $h$ (in colours, $s$). For visualization purposes, values on the y-axis are cut off at the threshold $t_c = 125$.

#### Appendix S9. Scaling of *s* and *u* by $2N_e$

The results presented in the main text relate to very small population sizes, of the magnitude of those used in laboratory experiments. The question of whether deleterious mutations can be purged after a much larger bottleneck, of the size more characteristic of natural populations, is also of interest. The present results would be relevant to this situation if the principle that the evolution of a finite population is determined by the products of twice the variance effective population size ($2N_e$) and the parameters that describe the intensity of the various evolutionary forces (selection coefficients, mutation rates, migration rates, recombination rates), when time is measured in units of $2N_e$ generations (the coalescence timescale). This approach is valid if the assumptions of diffusion equation theory are met (see Ewens 2004, Chapters 4 and 5), and has been widely used for reducing the computation time in simulations (Marsh et al. 2026; Johri et al. 2026).

If this principle were to hold in the present case, the results from the single-locus simulations and linear operator method could be extended to bottlenecked populations of size $CN$ instead of $N$, with $s$ and $u$ being replaced by $s/C$ and $u$/C, respectively, with time in generations being multiplied by $C$. Since the values of $\bar{q}$, $\bar{L}$ and $\bar{B}$ relative to their values for the ancestral population are not very sensitive to $u$, the rescaling would mainly concern $s$ and $N$. It faces the problem, however, that the initial expected frequency of a deleterious mutation in the bottlenecked population, conditioned on its surviving the bottleneck, is approximately equal to the product of the mean allele frequency in the initial population, $q^*$, and $2N$, which cannot be rescaled. In contrast, the dynamics of the probability distribution of $q$ after the bottleneck follows the standard diffusion equation, provided that the conditions for validity of the diffusion equation approximation hold.

A general solution for this probability distribution, based on the equivalent of Equation (3b) of the main text, has been obtained by Song and Steinrücken (2012), extending the previous work of Kimura (1955, 1957) for the cases of neutrality and semi-dominant selection. If the initial frequency of $A_2$, conditioned on its surviving the bottleneck is $q_0$, the probability density function at time $t$ for the post-bottleneck allele frequency conditioned on $A_2$ surviving the bottleneck takes the general form:

$$\phi(q, q_0, t) = \sum_{i=0}^{\infty} \psi_i\,(q_0)\varphi_i(q) e^{-\lambda_i t} \qquad \text{(S18)}$$

where $\psi_i$ and $\varphi_i$ are functions that correspond to the eigenvalue $\lambda_i$ ( $\lambda_i \geq 0$ for all $i$) (see Equation 5 of Song and Steinrücken 2012). These functions are complicated in form. However, $q_0$ is necessarily small (1/2*N*) in the present case, and $\psi_i(0) = 0$, so that Taylor's theorem implies that $\psi_i(q_0)$ is approximately proportional to 1/2*N*, provided that $\psi_i$ is a well-behaved function. The probability density for the unconditional frequency of $A_2$ is the product of $\phi(q, q_0, t)$ and $2Nq*$, so the factor of $2N$ cancels. Thus, rescaling should apply as a good approximation, provided that second-order terms in 1/(2*N*) and $spq[h + (1 - 2h)q]$ can be neglected in both the original and rescaled populations.

The accuracy of rescaling in the present case can be tested by comparing the results of the single-locus simulations and linear operator calculations for fixed values of the scaled selection coefficient but different population sizes. Figures S10 and S11 display some representative results, for cases with relatively strong and relatively weak selection, with $h$ = 0.2 and $h$ = 0.1, respectively. It can be seen that there is approximate agreement between the time courses (on the scale of 2*N* generations) of the relative values of $\bar{q}$, $\bar{F}$, $\bar{L}$ and $\bar{B}$ (relative to their values in the ancestral population) for a ten-fold difference in *N*. It thus seems reasonable to assume that our conclusions can be extrapolated to much larger sizes of bottlenecked populations. Agreement between the linear operator and simulation results is excellent for the weak selection cases shown in the right-hand panels of the two figures. It is also fairly close for the strong selection cases in the left-hand panels of Figure S10, for which $h$ = 0.2, but is substantially less good for the strong selection cases in the left-hand panels of Figure S11, where $h$ = 0.1. This probably reflects the fact that 2*Nhs* in this case is only 0.5, so that the assumptions used to obtain the linear operator results with strong selection are inaccurate. Note that the simulation values are means over replicate simulations, and are subject to significant error, accounting the vagaries of some of the curves in the figures.

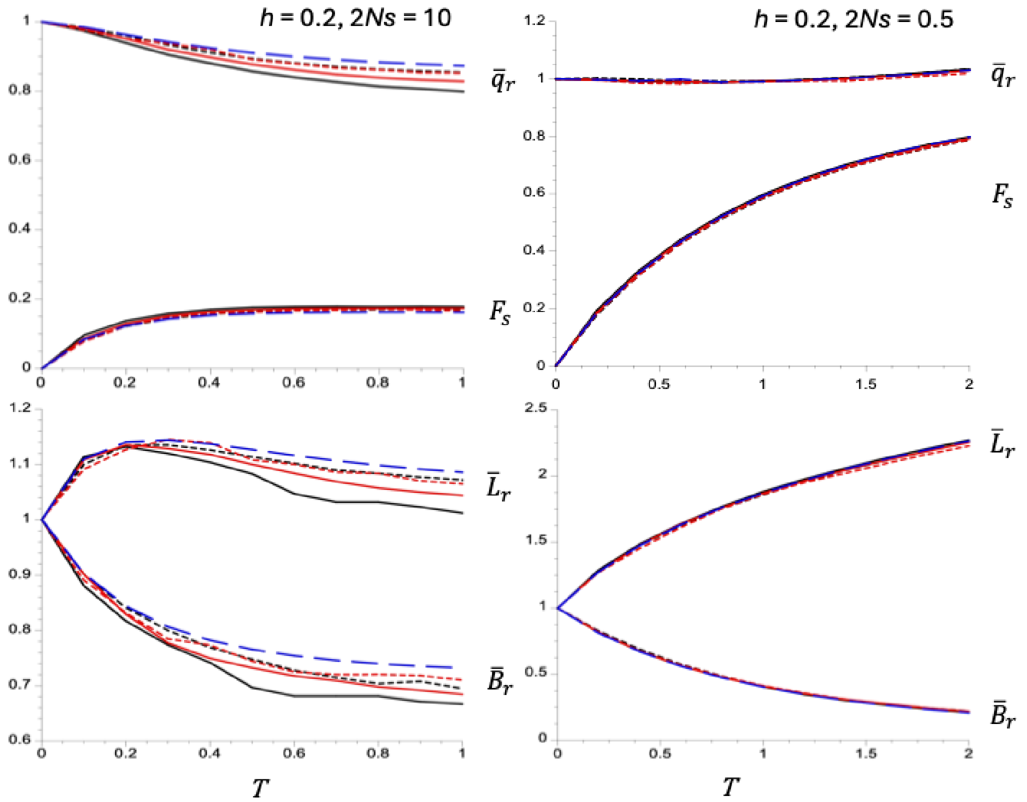


Figure S10. Plots against the time since the bottleneck in units of 2$N$ generations ($T$) of the mean frequency of $A_2$ relative to its ancestral value ($\bar{q}_r$), the fixation index at selected sites ($F_s$), the genetic load relative to its ancestral value ($\bar{L}_r$), and the inbreeding load relative to its ancestral value ($\bar{B}_r$).

For the left-hand panels (2$Ns$ =10, $h$ = 0.2), the solid black lines are single-locus simulation results for $N$ = 25, $s$ = 0.2, $u$ = 6.88 × $10^{-8}$ and $q*$ = 1.72 × $10^{-6}$; the dashed black lines are simulation results for $N$ = 250, $s$ = 0.02, $u$ = 6.88 × $10^{-9}$ and $q*$ = 1.72 × $10^{-6}$; the solid red lines are simulation results for $N$ = 50, $s$ = 0.1, $u$ = 7.03 × $10^{-8}$ and $q*$ = 3.51 × $10^{-6}$; the dashed red lines are simulation results for $N$ = 500, $s$ = 0.01, $u$ = 7.03 × $10^{-9}$ and $q*$ = 3.51 × $10^{-6}$; the broken blue lines are the results from linear operator Method 2a calculations for $N$ = 50, $s$ = 0.1, $u$ = 7.03 × $10^{-8}$ and $q*$= 3.51 × $10^{-6}$.

For the right-hand panels (2$Ns$ =0.5, $h$ = 0.2), the solid black lines are single-locus simulation results for $N$ = 25, $s$ = 0.01, $u$ = 7.76 × $10^{-9}$ and $q*$ = 3.88 × $10^{-6}$; the dashed black lines are simulation results for $N$ = 250, $s$ = 0.001, $u$ = 7.76 × $10^{-10}$ and $q*$ = 3.88 × $10^{-6}$; the solid red lines are simulation results for $N$ = 50, $s$ = 0.005, $u$ = 3.88 × $10^{-9}$ and $q*$ = 3.88 × $10^{-6}$; the dashed red lines are simulation results for $N$ = 500, $s$ = 0.0005, $u$ = 3.88 × $10^{-10}$ and $q*$= 3.88 × $10^{-6}$; the broken blue lines are the results from linear operator Method 2c calculations for $N$ = 50, $s$ = 0.005, $u$ = 3.88 × $10^{-9}$ and $q*$ = 3.88 × $10^{-6}$.

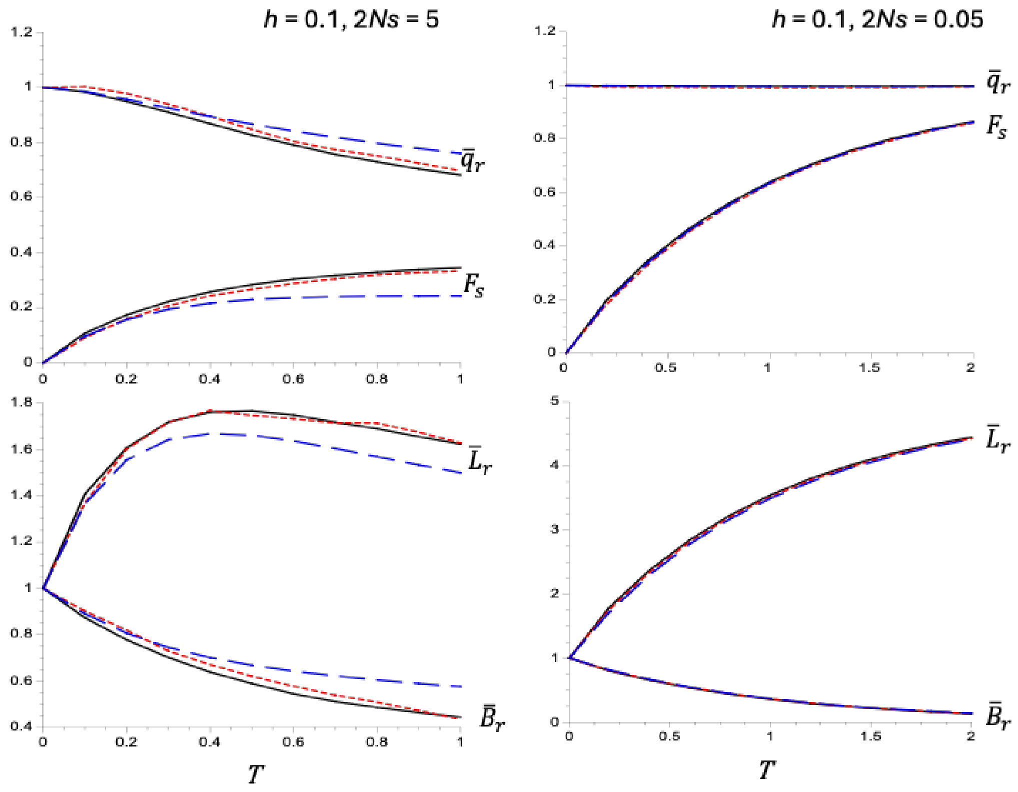


Figure S11. Plots against the time since the bottleneck in units of 2*N* generations (*T*) of the mean frequency of $A_2$ relative to its ancestral value ($\bar{q}_r$), the fixation index at selected sites ($F_s$), the genetic load relative to its ancestral value ($\bar{L}_r$), and the inbreeding load relative to its ancestral value ($\bar{B}_r$).

For the left-hand panels (2*Ns* = 5, *h* =0.1), the solid black lines are single-locus simulation results for *N* = 25, *s* = 0.1, *u* = 2.90 × $10^{-8}$ and *q** = 2.90 × $10^{-6}$; the dashed red lines are simulation results for *N* = 250, *s* = 0.01, *u* = 2.90 × $10^{-9}$ and *q** = 2.90 × $10^{-6}$; the broken blue lines are the results from linear operator Method 2a calculations for *N* = 50, *s* = 0.05, *u* =1.45 × $10^{-9}$ and *q** = 2.90 × $10^{-6}$.

For the right-hand panels (2*Ns* = 0.05, *h* =0.1), the solid black lines are single-locus simulation results for *N* = 25, *s* = 0.001, *u* = 4.23 × $10^{-9}$ and *q** = 4.23 × $10^{-5}$; the dashed red lines are simulation results for *N* = 250, *s* = 0.0001, *u* = 4.23 × $10^{-10}$ and *q** = 4.23 × $10^{-5}$; the broken blue lines are the results from linear operator Method 2c calculations for *N* = 50, *s* = 0.005, *u* = 2.11 × $10^{-9}$ and *q** = 4.23 × $10^{-5}$.

#### Appendix S10. Relation between $\bar{q}_r$ and the ratio of diversity at selected sites to diversity at neutral sites

The ratio of the expected value of $B$ for a given $h$ and $s$ to its value in the ancestral population is $\bar{B}_r = (1 - F_s)\bar{p}_r \, \bar{q}_r$, which is equal to the corresponding ratio of diversities at selected sites, $\pi_{sel\,r}$. The similar ratio for neutral sites, $\pi_{neut\,r}$, is equal to $1 - F$, where $F$ is the neutral fixation index. This implies that $\pi_{sel\,r}/\pi_{neut\,r}$ is given by:

$$\frac{\pi_{sel\,r}}{\pi_{neut\,r}} = \frac{(1-F_s)\bar{p}_r \, \bar{q}_r}{(1-F)} \equiv \frac{\bar{B}_r}{(1-F)} \qquad \text{(S19)}$$

In the rare-allele case, $\bar{p}_r$ can be approximated by 1, and the condition for $1 < \pi_{sel\,r}/\pi_{neut\,r}$ is simply:

$$\bar{q}_r > \frac{(1-F)}{(1-F_s)} \equiv \bar{B}_r > (1-F) \qquad \text{(S20)}$$

The same general conclusion applies if there is a heterogeneity in $s$ and $h$ among sites, as is the case in nature. For a given class $i$ of selected sites, there will be a value $F_{s\,i}$ of the fixation index and $\bar{q}_i$ of the mean allele frequency for the bottlenecked population at a given time. If expectations are taken across all sites, we have:

$$\mathrm{E}\{\bar{q}_i(1 - F_{s\,i})\} = \bar{q}(1 - \bar{F}_s) + \mathrm{Cov}\{\bar{q}_i(1 - F_{s\,i})\} \qquad \text{(S21)}$$

where $\bar{q}$ and $\bar{F}_s$ are the expected overall values of $\bar{q}_i$ and $F_{s\,i}$.

The covariance term is expected to be negative, since smaller values of $F_{s\,i}$ are associated with stronger selection, and hence smaller $\bar{q}_i$. Under the rare-allele assumption for selected sites, it follows that the inequality (S20) still holds if $F_s$ is replaced by $\bar{F}_s$. In general, therefore, it is easier to satisfy $1 < \pi_{sel\,r}/\pi_{neut\,r}$ than $\bar{q}_r > 1$.
